# Maladaptive immune-fibrotic axis drives impaired long bone regeneration under mechanical instability

**DOI:** 10.1101/2023.10.26.564177

**Authors:** Matthew D. Patrick, Jaimo Ahn, Kurt D. Hankenson, Ramkumar T. Annamalai

## Abstract

Delayed and non-healing fractures, affecting 5–10% of cases, are associated with prolonged disability and diminished quality of life. Although acute inflammation is required to initiate repair, persistent mechanical instability can sustain maladaptive immune and fibrotic responses that impede regeneration. Existing animal models do not adequately recapitulate mechanical instability, the principal driver of hypertrophic nonunion in clinical settings, thereby limiting translational relevance.

In this study, we developed a murine model of delayed fracture healing using tunable intramedullary fixation to impose controlled interfragmentary strain. High-strain conditions (low-stiffness nail, 15-30% strain) produced enlarged calluses characterized by delayed ossification, increased fibrotic tissue (2.9-fold, *p = 0.0099*), and reduced biomechanical integrity (1.6-fold decrease in stiffness*, p = 0.024*) relative to low-strain controls (high-stiffness nail, <5% strain). Spatial transcriptomic analysis identified persistent fibrotic niches in high-strain calluses enriched with fibroblast-associated genes (e.g., *Pdgfrb*, *Lgals3*) and dysregulated macrophage-fibroblast signaling (*Spp1*, *Mmp9*).

Systemically, high-strain fractures were associated with distinct immune signatures in which CD206⁺ macrophages and CD25⁺ regulatory T cells predicted healing outcomes (R = 0.72, *p = 0.004*), suggesting early immune polarization as a determinant of repair trajectory. Elevated CD8⁺ T cell responses were also observed, consistent with a sustained inflammatory state associated with impaired healing.

These findings identify mechanical instability as a driver of pathological immune-stromal interactions and establish a preclinical platform for investigating mechanobiology-informed therapeutic strategies. This work supports a conceptual framework in which hypertrophic nonunion is understood as a disorder arising from dysregulated interactions between mechanical cues and immune responses.

**Teaser:** Mechanical strain hijacks immune signaling to induce fibrosis and block bone regeneration in unstable fractures.

## A. Introduction

Fractures with delayed healing or nonunion remain a life-altering clinical challenge, affecting 5–10% of the 8 million annual fractures in the U.S., with profound impacts on patient morbidity and quality of life that can surpass chronic conditions like stroke, diabetes, and heart failure (*1–3*). Despite this impact, there is no standardized approach for early diagnosis, and confirmation of nonunion typically requires 6-9 months following injury or fixation (*4, 5*). This prolonged diagnostic window reflects an incomplete understanding of the biological and biomechanical processes that govern failed fracture repair (*3*), thereby limiting the identification of predictive biomarkers and targeted therapeutic strategies.

Among nonunions, the hypertrophic subtype is the most prevalent and is widely attributed to mechanical instability within the fracture environment (*6–8*). In contrast to atrophic nonunion, which is characterized by biological insufficiency, hypertrophic nonunion arises in settings where excessive interfragmentary strain disrupts the progression from fibrous and cartilaginous intermediates to mineralized bone (*7, 8*). Elevated strain has also been associated with amplification of inflammatory signaling pathways (*9, 10*). However, the mechanisms by which sustained mechanical instability redirects repair toward maladaptive outcomes remain insufficiently defined, in part because existing animal models do not adequately reproduce the strain-driven pathophysiology observed in clinical hypertrophic nonunion (*11*).

Bone possesses a distinctive capacity for scar-free regeneration under stable conditions (*12, 13*). Mechanical instability perturbs this regenerative sequence by maintaining elevated strain at the fracture site, thereby interfering with endochondral ossification (*14*). In delayed unions and nonunions, fibrotic tissue frequently accumulates within the fracture gap, forming a pseudarthrosis that compromises structural integrity and function (*2, 11*). Although fibrosis has traditionally been associated with atrophic nonunion in the context of segmental defects (*12*), clinical observations indicate that instability alone, even in the presence of bony apposition, can induce fibrotic healing responses (*3*). These observations suggest that fibrosis is not solely a consequence of biological insufficiency but may also arise from aberrant mechanical environments.

Strain theory provides a quantitative framework for understanding tissue differentiation during fracture repair, positing that local strain magnitude determines the predominant tissue phenotype (*15–17*). Empirical and computational studies indicate that interfragmentary strains exceeding approximately 15% favor fibrous tissue formation, whereas strains below 5% support bone formation (*18*). During normal healing, progressive increases in callus stiffness reduce local strain, enabling the transition to ossification (*19*). In hypertrophic nonunion, this transition fails to occur, and the callus remains in a persistent high-strain state, typically within the range of 15-30%, which promotes fibrotic tissue persistence and prevents bridging bone formation. This mechanical environment is likely to influence not only stromal cell behavior but also immune cell function, thereby shaping the overall trajectory of repair. However, conventional nonunion models, which rely on critical-size defects or the absence of fixation, do not isolate strain as a primary variable and therefore provide limited insight into strain-mediated mechanisms (*11, 20*).

Within the fracture microenvironment, macrophages and fibroblasts are positioned as central regulators of repair and fibrosis. Mechanical strain can modulate cellular behavior through mechanotransductive pathways, including focal adhesion signaling and tension-mediated activation of latent transforming growth factor beta 1 (TGF-β1), a key driver of myofibroblast differentiation (*21*). Syndecan-4, acting in concert with integrins, functions as a mechanosensitive co-receptor that regulates downstream pathways involved in extracellular matrix remodeling (*22, 23*). These signaling cascades converge on profibrotic mediators such as TGF-β, osteopontin (SPP1), and galectin-3, which promote matrix deposition and tissue remodeling across multiple organ systems (*24, 25*). In addition, thrombospondins can activate latent TGF-β and may act synergistically with syndecan-4–dependent pathways to reinforce fibrosis under conditions of mechanical stress (*26*). While these mechanisms are well described in other fibrotic contexts, it remains uncertain whether elevated mechanical strain alone is sufficient to initiate and sustain fibrotic remodeling during fracture healing *in vivo*.

Fracture repair is further influenced by systemic immune responses that can modulate regenerative capacity (*10, 20*). Prior studies indicate that early systemic immune profiles are associated with subsequent healing outcomes and that mechanical instability can alter these responses (*27*). Imbalances in circulating CD8⁺ T cell subsets and regulatory T cells (Tregs) have been linked to impaired healing, while Tregs have been shown to support bone regeneration (*27–29*). Additionally, CD206⁺ macrophages have been observed in fibrotic and poorly healing callus tissue (*28–30*). These findings suggest that strain-induced alterations in the local microenvironment may be reflected systemically through distinct immune signatures, providing a potential basis for biomarker development.

We hypothesized that sustained high interfragmentary strain is sufficient to maintain a profibrotic healing state through altered macrophage–fibroblast signaling, whereas low-strain conditions permit progression toward ossification and functional repair. We further hypothesized that this strain-dependent divergence in healing outcomes would be reflected in circulating immune phenotypes associated with delayed healing. To test these hypotheses, we adapted a blunt-trauma fracture model (*31, 32*) and developed a tunable intramedullary fixation system using nitinol (nickel–titanium) implants to generate controlled low-strain (<5%) and high-strain (15–30%) environments by varying nail stiffness. This approach enabled isolation of mechanical instability as the primary experimental variable. Using this platform, we examined the effects of strain on fibrotic fracture healing, characterized associated immune-stromal signaling pathways, including thrombospondin (*Thbs1*/TSP1)–syndecan-4 interactions and galectin-3–associated macrophage (*Lgals3^+^*) responses, and evaluated corresponding systemic immune signatures, including CD206⁺ macrophages, regulatory T cells, and CD8⁺ T cells, as candidate biomarkers of delayed healing and nonunion risk.

## B. Results

### 1. Modulating Interfragmentary Strain via Customized Intramedullary Nail Stiffness

To investigate the impact of mechanical instability on fracture healing, we engineered intramedullary (IM) nails from superelastic nitinol (NiTi) with defined stiffness profiles (3–30 N/mm, **Supplementary Fig. S1**). Three-point bending tests identified two configurations that generated distinct interfragmentary strain environments under physiological loading: a low-stiffness nail (2.7 ± 0.05 N/mm) designed to permit high strain (>15%), and a high-stiffness nail (16.5 ± 0.41 N/mm) that limited strain to <5% (**Fig. 1A**). Prior work demonstrated that medical-grade 316L stainless-steel nails (44 ± 3.3 N/mm; diameter 0.51 mm) produce healing outcomes comparable to the high-stiffness configuration; accordingly, the high-stiffness group was used as the control condition.

**Figure 1.**
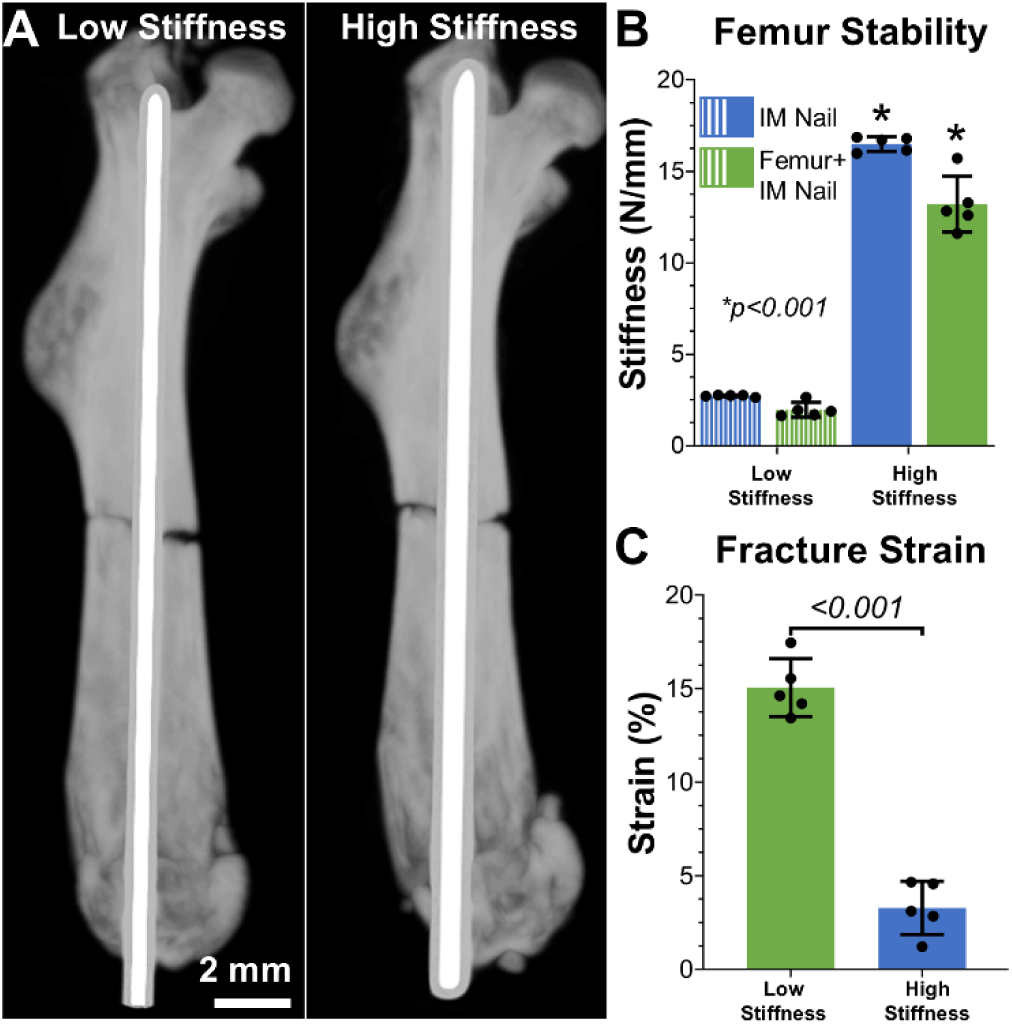
Intramedullary Nail Stiffness Modulates Interfragmentary Strain During Fracture Healing. **(A)** Representative maximum-intensity projection microCT scans of murine femurs stabilized with low- or high-stiffness intramedullary (IM) nails. **(B)** Stiffness of standalone nitinol nails versus nails implanted in fractured femurs (n = 5 per group). **(C)** Flexural strain at the fracture site under physiological loads (n = 5 per group; mean ± SD).

Biomechanical testing revealed that stabilized femurs exhibited slightly reduced stiffness compared to standalone nails (low-stiffness group: 1.96 ± 0.41 N/mm vs. nail alone: 2.7 N/mm; high-stiffness group: 13.21 ± 1.53 N/mm vs. nail alone: 16.5 N/mm; **Fig. 1B**), consistent with load sharing between implant and bone. Under physiological loading conditions (1.4–2.1 N, approximating murine hindlimb forces), the low-stiffness configuration generated interfragmentary strains of 15.1 ± 1.4%, whereas the high-stiffness configuration produced strains of 3.3 ± 1.4% (**Fig. 1C**). These values correspond to established thresholds associated with fibrous tissue formation (>15%) and bone formation (<5%), respectively. Together, these results validate a fixation platform that reproducibly generates distinct mechanical environments, thereby enabling controlled investigation of strain-dependent fracture-healing responses.

### 2. Interfragmentary Strain Governs Callus Morphology and Composition

To determine how mechanical strain influences fracture healing outcomes, transverse femoral fractures in 22-week-old mice (n = 11 per group) were stabilized using either high-stiffness (∼15 N/mm) or low-stiffness (∼2 N/mm) intramedullary nails. Analyses were conducted at 3 weeks post-fracture, corresponding to a stage of hard callus formation with ongoing remodeling in murine femoral repair (*33*).

MicroCT analysis demonstrated marked differences in callus morphology between groups. High-strain fractures (low-stiffness nail, >15% interfragmentary strain) developed substantially larger calluses, with a 1.7-fold increase in total tissue volume (TV, *p = 0.003*) and a 1.6-fold increase in bone volume (BV, *p < 0.001*) relative to low-strain controls (high-stiffness nail, <5% strain) (**Fig. 2A–C, 2D–E**; **Table 1**). Total mineral content was also elevated in the high-strain group (1.8-fold increase, *p < 0.001*; **Fig. 2F**), consistent with morphological features reported in hypertrophic nonunion (16).

**Figure 2.**
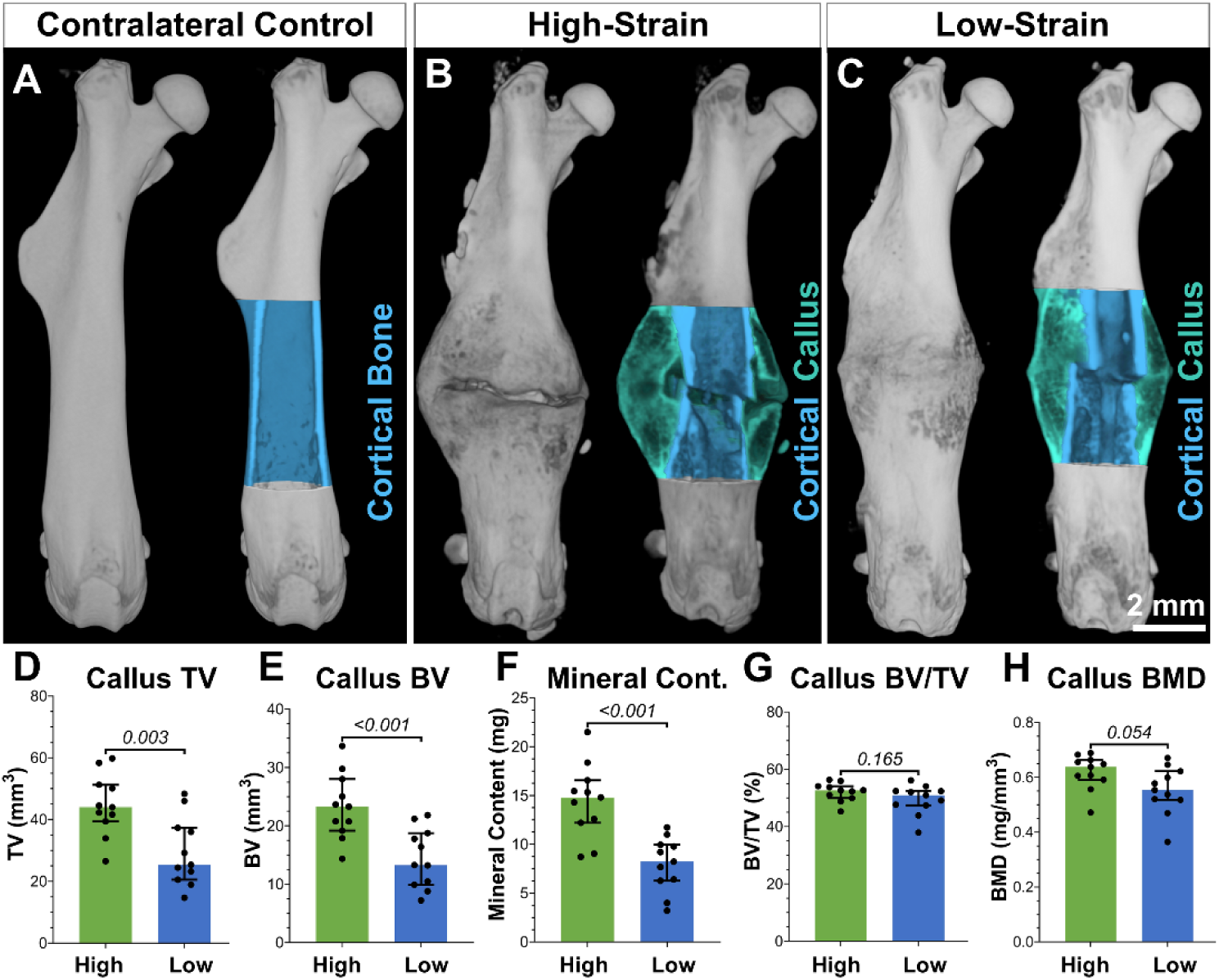
Interfragmentary Strain Dictates Callus Morphology and Composition at 3 Weeks Post-Fracture. (**A**) Representative 3D microCT reconstructions of contralateral uninjured femurs (control). (B) High-strain group (low-stiffness nail, >15% interfragmentary strain) and (C) Low-strain group (high-stiffness nail <5% interfragmentary strain), with transverse cross-sections highlighting the fracture site (volume of interest: ±2.5 mm from the fracture line). Cortical bone (light blue) and callus tissue (teal) are demarcated. (D–H) Quantitative analysis of callus tissue volume (TV), bone volume (BV), mineral content (MC), bone volume to tissue volume ratio (BV/TV), and bone mineral density (BMD) in high-strain vs. low-strain groups (n = 11 per group; mean ± SD; significance defined as p < 0.05, unpaired t-test).

**Table 1.** Bone morphometric analysis. Comparative evaluation of bone morphometric properties between the high-strain (low-stiffness nail) and low-strain (high-stiffness nail) groups. The values presented are expressed as mean±SD. n = 11 per group.

| **Morphometric properties** | **High-Strain** | **Low-Strain** | **p-value** |
| --- | --- | --- | --- |
| Callus TV (mm<sup>3</sup>) | 44.72 ± 9.9 | 29.47 ± 11.1 | 0.0028 |
| Callus BV (mm<sup>3</sup>) | 23.31 ± 5.6 | 14.45 ± 5.1 | 0.0009 |
| Callus Mineral content (mg) | 14.45 ± 3.8 | 7.95 ± 2.8 | 0.0002 |
| Callus BV/TV (%) | 52.08 ± 3.2 | 49.45 ± 5.1 | 0.1652 |
| Callus BMD (mg/mm<sup>3</sup>) | 0.62 ± 0.1 | 0.55 ± 0.1 | 0.0536 |
| Callus Trabecular Thickness (mm) | 0.115 ± 0.003 | 0.117 ± 0.004 | 0.2391 |
| Callus Trabecular Number (mm<sup>-1</sup>) | 4.51 ± 0.3 | 4.23 ± 0.5 | 0.140 |
| Callus Trabecular Separation (mm) | 0.16 ± 0.04 | 0.16 ± 0.1 | 0.8180 |
| Callus Bone Surface (mm<sup>2</sup>) | 618.89 ± 140.6 | 384.03 ± 129.1 | 0.0006 |
| Callus Bone Surface Density (mm<sup>2</sup>/mm<sup>3</sup>) | 13.85 ± 0.7 | 12.98 ± 1.4 | 0.0902 |
| Callus Specific bone surface (mm<sup>2</sup>/mm<sup>3</sup>) | 26.61 ± 0.9 | 26.78 ± 1.1 | 0.6916 |
| Callus Structure Model Index (SMI) | -0.69 ± 0.3 | -0.26 ± 0.5 | 0.0204 |
| Callus Connectivity density (mm<sup>-3</sup>) | 196.87 ± 23.1 | 167.57 ± 29.5 | 0.0174 |
| Callus Degree of anisotropy | 1.25 ± 0.1 | 1.43 ± 0.2 | 0.0054 |

In contrast, bone volume fraction (BV/TV) and bone mineral density (BMD) did not differ significantly between groups (**Fig. 2G–H**), indicating that while mechanical instability promotes callus expansion, it does not measurably alter mineralization efficiency at this stage of healing.

Histological analysis further delineated strain-dependent healing mechanisms. Movat’s pentachrome staining of week-3 femurs revealed that high-strain calluses were dominated by vascularized fibrotic tissue (collagen: yellow; fibrin/muscle: red) with minimal chondrogenic regions (mucins: blue), while low-strain calluses exhibited mature woven bone with robust collagen deposition (**Fig. 3A(i–ii**), 3**B(i–ii)**). Immunohistochemical analysis identified regions of chondrogenesis (SOX9+), osteoblast activity (SP7+), vascularization (CD31+), and myofibroblast activation (α-SMA+) within both high-strain and low-strain fracture calluses (**Fig. 3A(iii–vi), 3B(iii–vi)**). Quantification revealed comparable levels of SOX9+ chondrogenic cells between strain groups across both the intramedullary (IM) and periosteal (PO) callus regions (**Fig. 3C(i)**). In contrast, SP7+ osteoblasts were higher in the low-strain group in both IM and PO compartments (**Fig. 3C(ii)**). CD31+ endothelial staining indicated similar levels of vascularization in the PO callus, although a downward trend was observed in the IM region of the high-strain condition (**Fig. 3C(iii)**). Notably, α-SMA+ myofibroblast accumulation was increased in the IM region of high-strain calluses, with comparable levels in the PO region across groups (**Fig. 3C(iv)**), suggesting localized fibrotic remodeling under elevated mechanical load.

**Figure 3.**
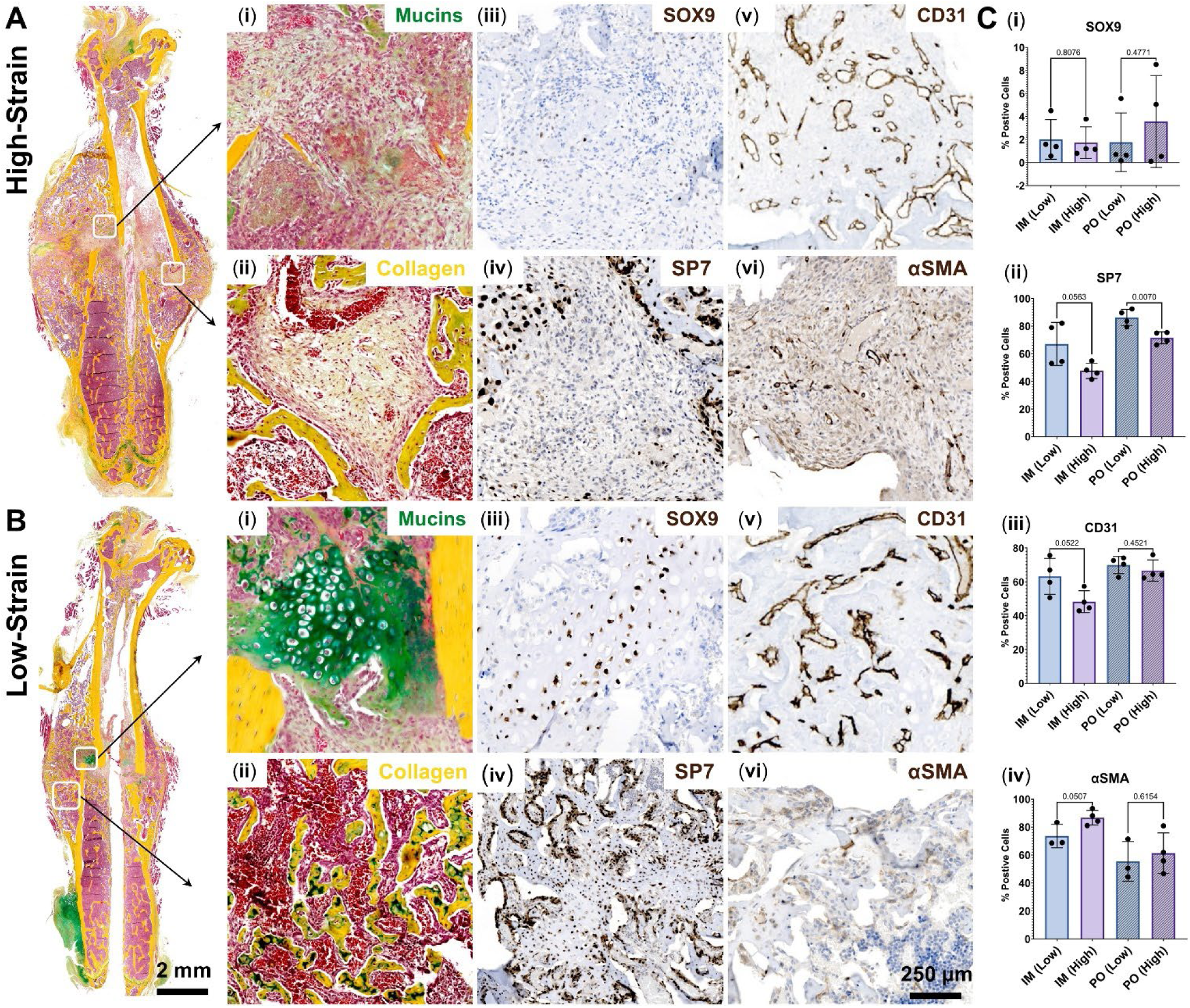
Interfragmentary Strain Modulates Endochondral Ossification and Fibrotic Remodeling. (**A**) High-strain (low-stiffness nail, >15% interfragmentary strain) and (**B**) Low-strain (high-stiffness nail, <5% interfragmentary strain) groups at 3 weeks post-fracture. Representative Movat’s pentachrome staining (i–ii) and immunohistochemistry (IHC) of fracture calluses. (i) High-strain callus showing fibrotic tissue (yellow collagen, red fibrin) and reduced chondrogenic areas (blue mucins). (ii) Low-strain callus with mature woven bone (yellow collagen) and minimal fibrosis. (iii) SOX9+ chondrocytes (brown) and (iv) SP7+ osteoblasts (brown) highlight lineage-specific activity. (v) CD31/PECAM+ endothelial cells (brown) indicate vascularization. (vi) α-SMA+ fibrotic regions (brown) dominate high-strain calluses. (**C**) (i–iv) Quantification of IHC as a percentage of positive cells in the intramedullary (IM) and periosteal (PO) callus under high-strain and low-strain conditions (n = 3–4 per group; mean ± SD).

Picrosirius red staining combined with polarized light imaging revealed an abundance of collagen-rich tissue areas within high-strain calluses compared to low-strain controls (**Fig. 4A(i–iii), B(i–iii)**). Spatial transcriptomics localized enhanced *Col3a1* expression to fibrotic regions of the high-strain calluses (**Fig. 4A(iv), B(iv)**), corroborating the molecular signature of pathological ECM remodeling. Quantification of the collagen tissue and architecture between groups revealed a 3.3-fold increase in collagen-rich tissue areas (p = 0.0147) in high-strain calluses along with an elevated total fiber count (p = 0.0721) (**Fig. 4C**), while the average fiber width (low-strain: 1.22 ± 0.05 µm, high-strain: 1.20 ± 0.06 µm), length (low-strain: 11.52 ± 0.74 µm, high-strain: 11.88 ± 0.57 µm), and angle (low-strain: 91.06 ± 5.81 θ, high-strain: 92.38 ± 8.65 θ) remained unchanged between these areas. Together, these findings establish that elevated interfragmentary strain drives collagen dysregulation and fibrosis, impairing functional bone healing, while mechanical stability promotes ossification and regenerative repair.

**Figure 4.**
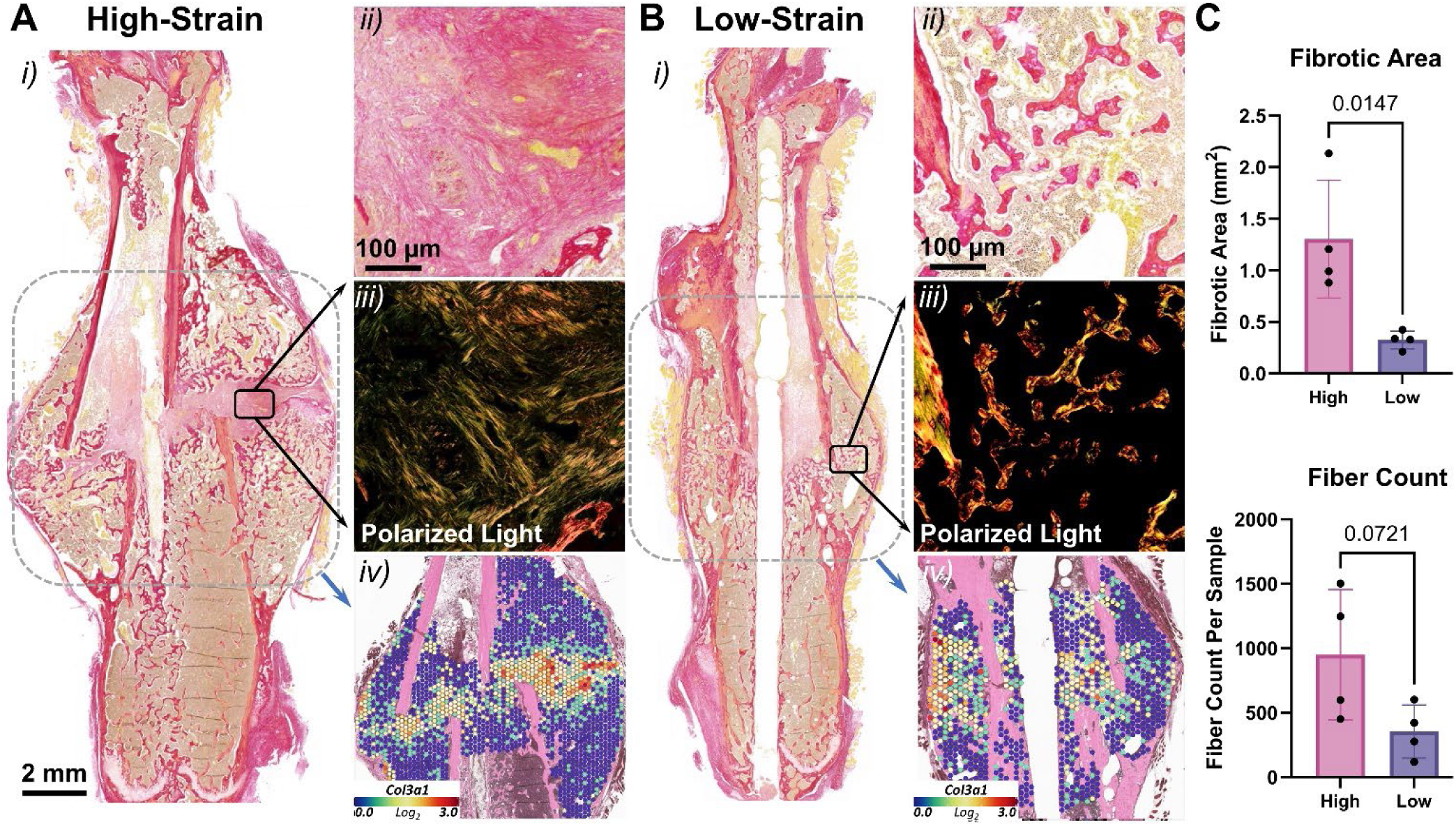
Interfragmentary Strain Drives Fibrotic Remodeling Through Collagen Reorganization and Col3a1 Expression. **(A)** High-strain (low-stiffness nail, >15% interfragmentary strain) and **(B)** Low-strain (high-stiffness nail, <5% interfragmentary strain) groups at 3 weeks post-fracture. **(i)** Picrosirius red staining (brightfield) identifies fibrotic collagen-rich regions (red). **(ii)** High-magnification views highlight dense collagen deposition. **(iii)** Polarized light imaging reveals parallel-aligned fibrillar collagen (bright birefringence) in high-strain calluses, indicative of mature fibrosis, versus disorganized collagen in low-strain groups. **(iv)** Spatial transcriptomics maps show elevated Col3a1 expression (red-purple) in fibrotic regions of high-strain calluses. Scale bars: 2 mm (i), 100 µm (ii–iii). (**C**) Quantification of fibrotic area (top) and total fiber count (bottom), showing elevated level of fibrotic tissue within the high-strain calluses (n = 4 per group; mean ± SD).

### 3. Mechanical Strain Activates Fibrogenic Pathways Through Transcriptional Reprogramming and Cellular Crosstalk

Spatial transcriptomics of week-3 calluses revealed distinct cellular niches shaped by interfragmentary strain (**Fig. 5A**). A representative high-strain and low-strain callus was analyzed using Visium (FFPE; 55 µm spot resolution). Fibroblast-enriched and osteoblast-enriched regions identified by transcriptomic clustering co-localized with picrosirius red–positive collagen and Osterix IHC on adjacent sections, supporting the anatomical and biological specificity of these domains.

**Figure 5.**
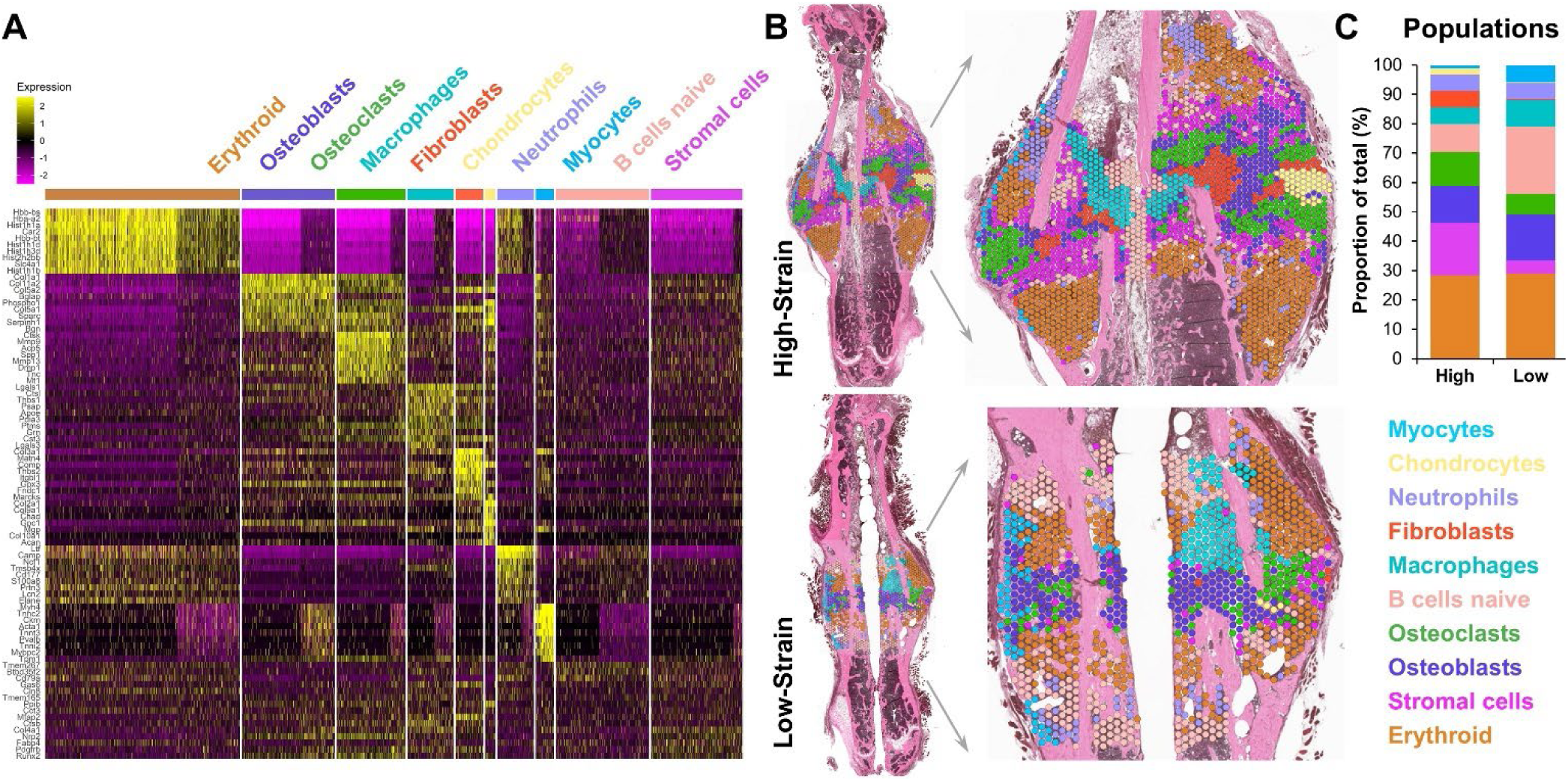
Mechanical Strain Induces Macrophage-Fibroblast Co-Localization in Fibrotic Callus Niches. (A) Heatmap of the top 10 differentially expressed genes across spatial transcriptomics clusters. (B) Histological sections of high-strain (low-stiffness nail, >15% interfragmentary strain, top) and low-strain (high-stiffness nail, <5% interfragmentary strain, bottom) calluses overlaid with spatial transcriptomics clusters (STclusters). Fibroblast-rich regions (STcluster, Red) in high-strain calluses overlap with picrosirius red-positive fibrotic zones (Fig. 4A). (C) Cellular composition analysis reveals a 14-fold increase in fibroblast-rich areas (5.79% vs. 0.39%) and 1.7-fold more osteoclast-rich regions (11.58% vs. 6.96%) in high-strain versus low-strain calluses. One representative FFPE callus from each group was used for spatial profiling; serial adjacent sections were processed for picrosirius red, Movat’s pentachrome, and IHC for orthogonal validation.

Unsupervised clustering of gene expression profiles identified fibroblast-rich regions comprising 5.79% of the high-strain callus, a 14-fold increase over the low-strain control (0.39%; **Fig. 5B**). These fibroblast-dense zones spatially overlapped with picrosirius red-positive fibrillar collagen deposits (**Fig. 4A**), confirming their fibrotic identity. The high-strain callus also exhibited a 1.7-fold elevation in osteoclast-rich areas (11.58% vs. 6.96% in low-strain, **Fig. 5C**), with osteoclasts localizing adjacent to fibroblast clusters, suggesting coupled remodeling of fibrotic and mineralized matrices.

Despite analogous spatial positioning of cellular niches in high-strain and low-strain calluses (**Fig. 6A**), transcriptional profiles diverged markedly. Fibrosis-associated genes (*Mmp9, Spp1, Pdgfrb, Lgals3*) mirrored patterns similar to those reported in pulmonary and hepatic fibrosis (**Fig. 6B**). Osteoclasts in the high-strain condition exhibited upregulated *Mmp9* (1.03-fold, p < 0.0001), a collagenase linked to pathological matrix degradation, while macrophage-rich regions showed elevated *Spp1* (osteopontin, 0.93-fold), *Pdgfrb* (0.75-fold), and *Lgals3* (galectin-3, 0.58-fold; *p < 0.0137*), key mediators of fibroblast activation and ECM stiffening. Macrophage-fibroblast proximity in the high-strain callus (**Fig. 6B**) suggested paracrine crosstalk, further supported by UMAP clustering, which revealed distinct fibrotic transcriptional signatures (**Fig. 6C–D**).

**Figure 6.**
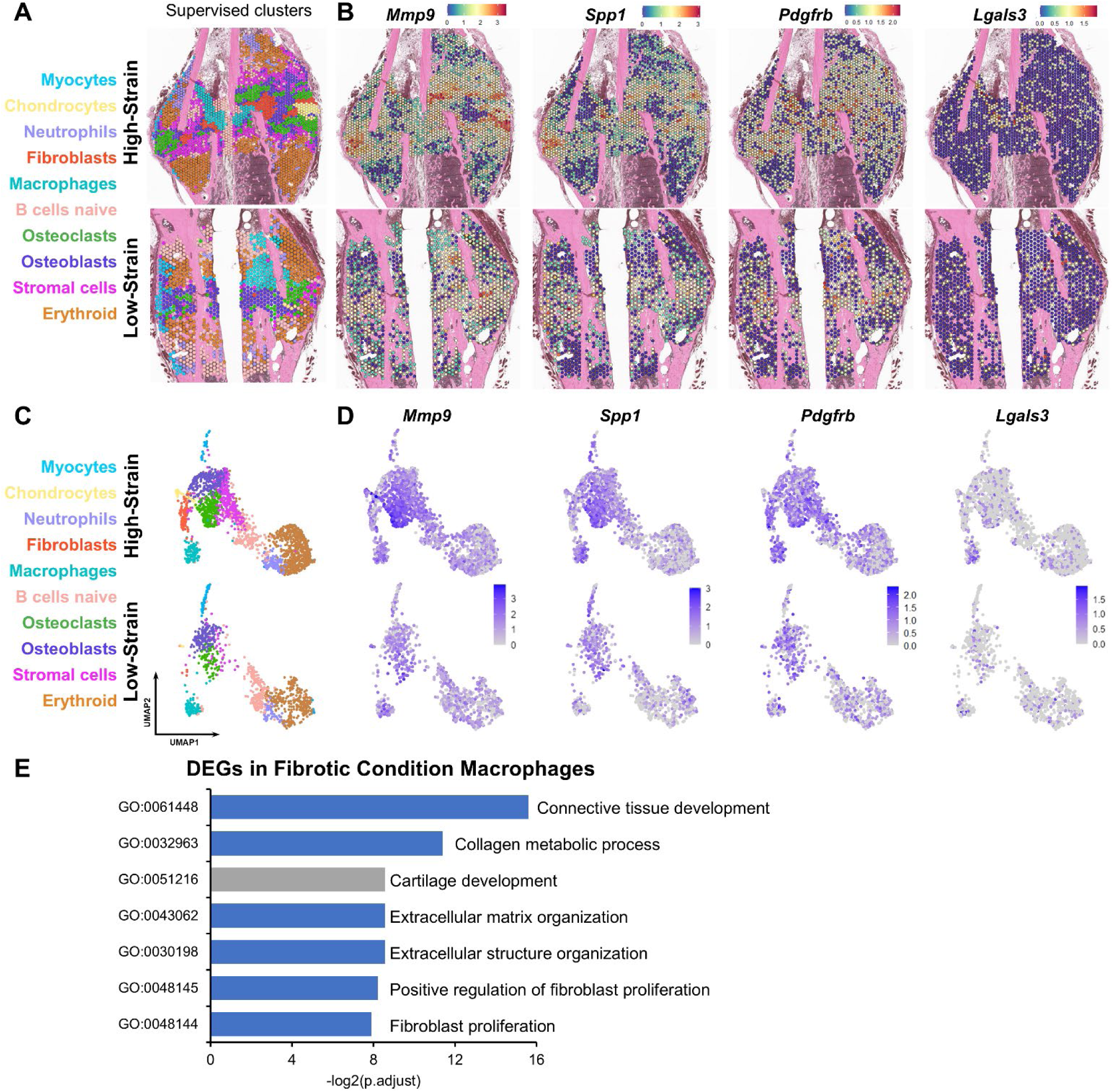
Strain-Dependent Transcriptional Reprogramming Drives Fibrotic Remodeling. (**A**) Histological sections of high-strain (low-stiffness nail, >15% interfragmentary strain, top) and low-strain (high-stiffness nail, <5% interfragmentary strain, bottom) calluses with overlaid spatial transcriptomics clusters. (**B**) Spatial expression maps of fibrosis-associated genes (Mmp9, Spp1, Pdgfrb, Lgals3) in high-strain and low-strain samples. (**C**) UMAP plots of cell clusters colored by type. (**D**) UMAP projections showing expression levels of fibrotic genes, with intensity representing expression levels. (**E**) Gene Ontology (GO) analysis of biological processes enriched due to differentially upregulated genes in macrophages in the high-strain condition. One representative FFPE callus from each group was used for spatial profiling; serial adjacent sections were processed for picrosirius red, Movat’s pentachrome, and IHC for orthogonal validation.

Analysis of cell-type-enriched areas within different callus compartments (**Fig. S2A**) revealed pronounced differences between the high-strain and low-strain fractures, mirroring the overall pattern observed by immunostaining. Chondrocyte-enriched regions were more prevalent in the PO of the high-strain fracture (6.15% of total PO) compared to the low-strain fracture (0.52% of total PO), a difference not evident in the immunostains. Osteoblast-enriched regions were expanded in the IM of the low-strain fracture, with a 3.54-fold increase (21.14% of total IM) relative to the high-strain fracture (5.96% of total IM), consistent with the significant differences observed in Sp7. Endothelial cell-enriched regions were not detected in either group. By contrast, fibroblast-enriched regions showed an 11.92-fold increase in the PO of the high-strain fracture (6.17% of total PO) compared to the low-strain fracture (0.52% of total PO), aligning with αSMA protein localization. To further compare with the immunostains in **Fig. 3**, we evaluated Sox9 (SOX9), Sp7 (SP7/Osterix), Pecam1 (CD31), and Acta2 (αSMA) expression in Visium sections (**Fig. S2B**). Sox9, Pecam1, and Acta2 showed low transcript abundance (as expected), whereas Sp7 was readily detected and recapitulated IHC localization but did not differ significantly between the high-strain and low-strain fractures across cell types or within PO or IM compartments.

Gene Ontology (GO) enrichment analysis of differentially expressed genes highlighted pathways central to fibroblast proliferation (*PDGF* signaling) and ECM organization (collagen fibril assembly), with macrophages as primary contributors (**Fig. 6E, Tables S1–S2**). Osteoclasts and osteoblasts showed enrichment in ECM disassembly and ossification pathways, respectively, underscoring their dual roles in matrix remodeling. Together, these findings position mechanical strain as a catalyst for maladaptive immune-stromal interactions that sustain fibrosis and impede functional repair.

CellChat analysis revealed enhanced communication between fibroblasts, macrophages, and osteoclasts in the high-strain callus. The high-strain group exhibited globally elevated cellular communication compared to the low-strain control, with fibroblasts, macrophages, and osteoclasts emerging as hubs for incoming and outgoing signals (**Fig. 7A**). Pathway analysis identified 16 mechano-sensitive pathways, including *TGFβ, ncWNT, PDGF, THBS, FN1, SPP1, TENASCIN*, and *COLLAGEN*. Notably, the *TENASCIN* and *THBS* pathways exhibited the most pronounced activation with the greatest differential signaling between strain conditions (**Fig. 7B**). Macrophages in the high-strain callus communicated via thrombospondin1/2 (*Thbs1/2)* and *Comp* ligands targeting fibroblast *Sdc4* receptors, while osteoclasts utilized *Tnc* and *Comp* to engage fibroblast receptors (**Fig. 7C–D**). These interactions were absent in the low-strain callus, underscoring strain-dependent modulation of cellular crosstalk.

**Figure 7.**
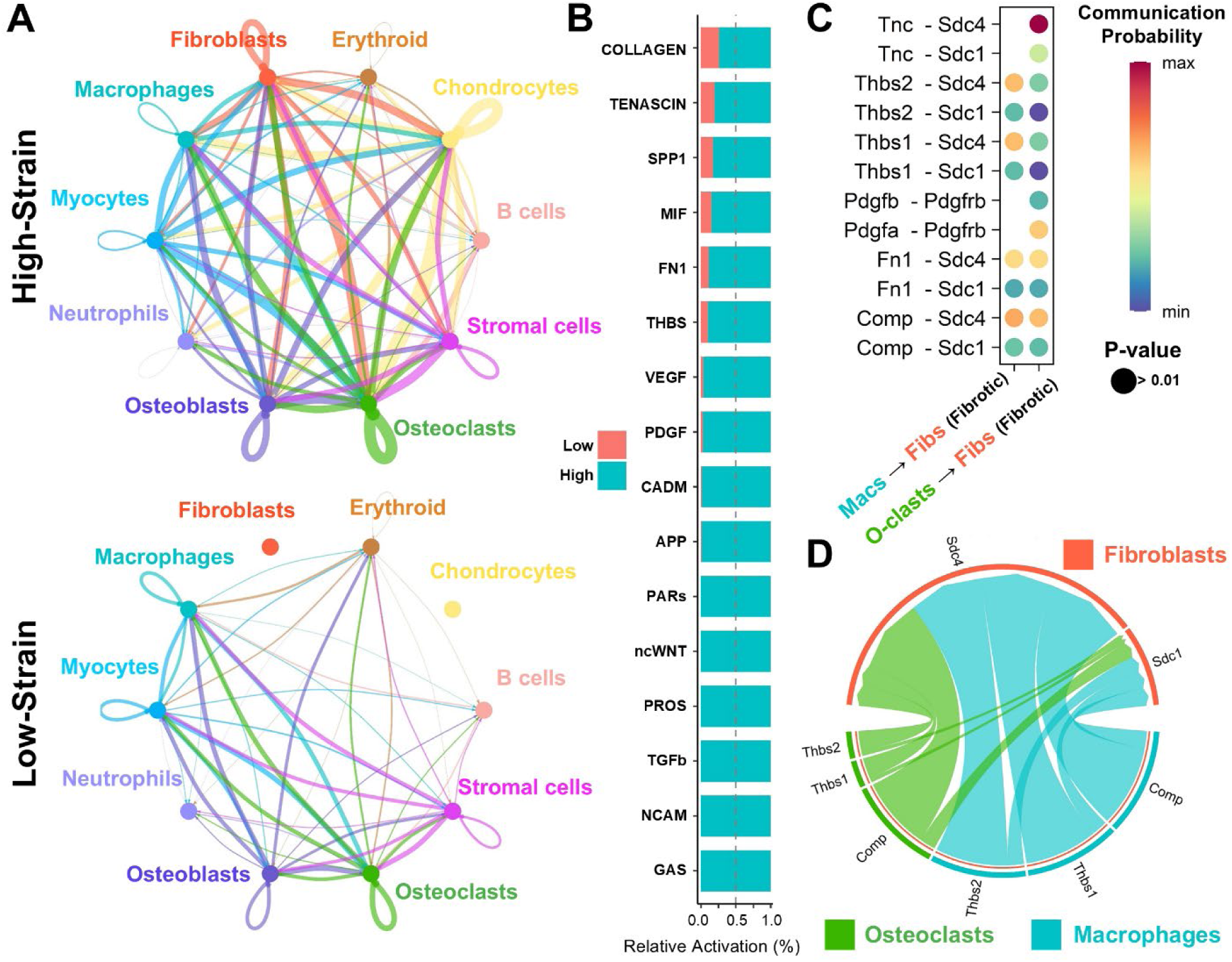
Mechano-Sensitive Cellular Networks Sustain Fibrotic Crosstalk. **(A)** CellChat interaction networks in high-strain (low-stiffness nail, >15% interfragmentary strain, top) and low-strain (high-stiffness nail, <5% interfragmentary strain, bottom) calluses. Line thickness denotes interaction strength. **(B)** Relative activation levels of key signaling pathways, with most pathways showing increased activity in the high-strain callus. **(C)** Communication probability scores for key ligand-receptor interactions, with higher probability indicated by warmer colors. **(D)** Chord diagram highlighting major signaling interactions in the high-strain callus, with fibroblasts (red), macrophages (green), and osteoclasts (blue) as key mediators. One representative FFPE callus from each group was used for spatial profiling; serial adjacent sections were processed for picrosirius red, Movat’s pentachrome, and IHC for orthogonal validation.

Guided by enrichment of THBS signaling between macrophages and fibroblasts (**Fig. 7**), we examined *Thbs1*/TSP1. Visium showed that *Thbs1* expression was concentrated along the IM fracture gap in the high-strain callus (**Fig. 8A**). On serial sections, TSP1 IHC localized to matrix/pericellular regions within the IM canal (**Fig. 8B**), colocalizing with macrophage- and fibroblast-enriched niches in high-strain calluses (**Fig. 5**). Quantification demonstrated increased TSP1-positive area in the IM compartment of high-strain calluses (*p = 0.0739*), with a weaker corresponding trend in the PO compartment (*p = 0.2006*) (**Fig. 8C**). Collectively, these data implicate *Thbs1*/TSP1-mediated macrophage-fibroblast crosstalk in strain-related fibrosis.

**Figure 8.**
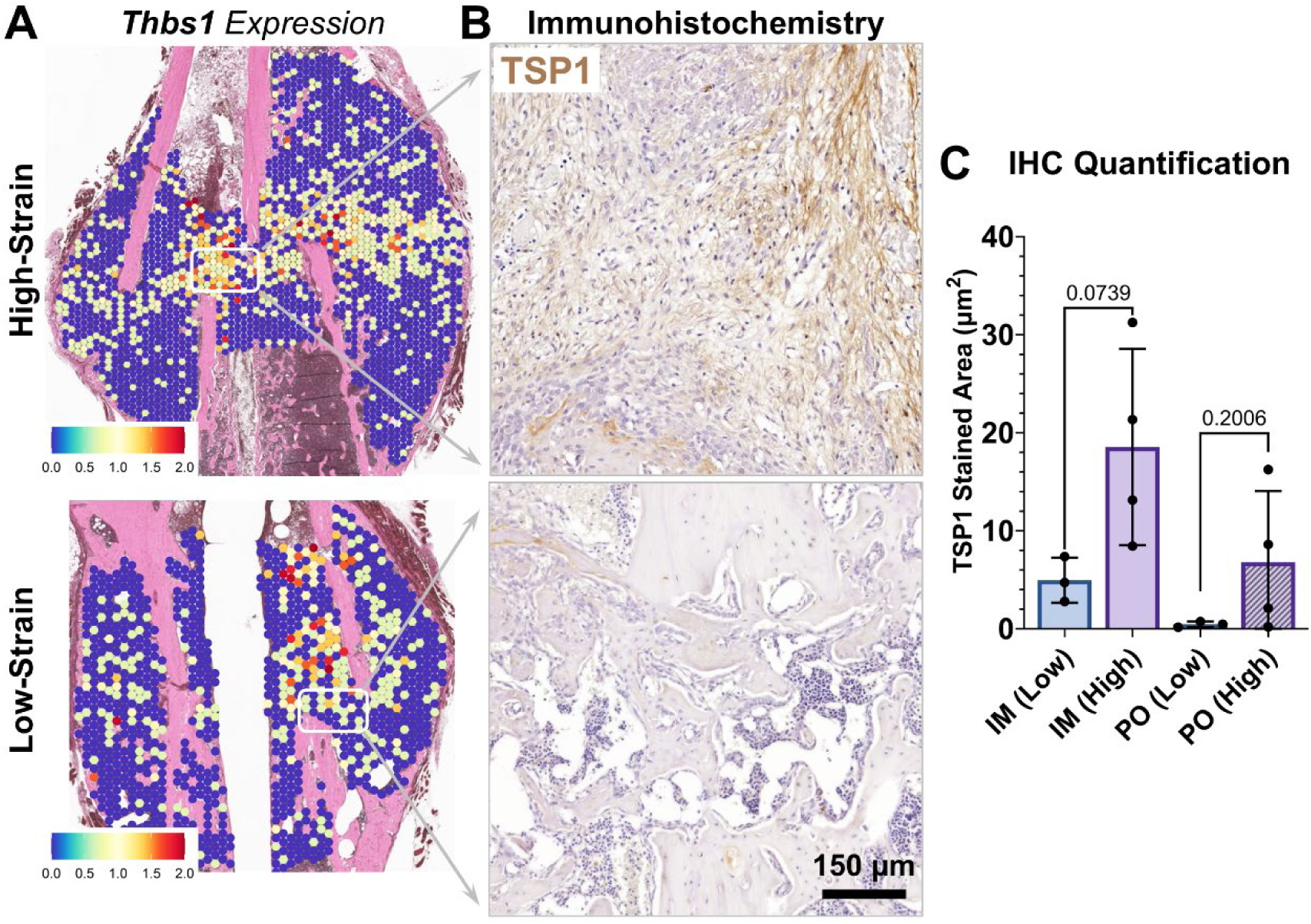
Thbs1 is Prevalent in High-Strain Fractures. **(A)** Spatial expression maps of Thbs1 in high-strain (low-stiffness nail, >15% interfragmentary strain, top) and low-strain (high-stiffness nail, <5% interfragmentary strain, bottom) groups. **(B)** TSP1 IHC on serial sections adjacent to the Visium slides shows matrix/pericellular staining enriched in high-strain callus (scale bar, 100 µm). **(C)** Quantification of TSP1-positive DAB area in intramedullary (IM) and periosteal (PO) compartments (n = 3–4 animals per group; mean ± SD; two-sided unpaired t-test.

### 4. Mechanical Instability Compromises Functional Healing Outcomes

Functional assessment by three-point bending at 3 weeks post-fracture revealed marked mechanical deficiencies in high-strain calluses (see **Table 2** for detailed values). Low-strain fractures exhibited superior whole-bone properties (**Fig. 9A–D)**, with 1.6-fold higher stiffness (*p* = 0.024, **Fig. 9A**), 1.4-fold greater maximum force (*p* = 0.036, **Fig. 9B**), and 1.7-fold elevated ultimate stress (*p* = 0.002, **Fig. 9G**) compared to high-strain fractures. Tissue-level properties (**Fig. 9E–H**) further demonstrated impaired healing in high-strain calluses, with yield stress reduced by 1.9-fold (*p* = 0.032, **Fig. 9H**). Force-at-yield, post-yield displacement (PYD), work-to-fracture, and elastic modulus did not differ significantly between groups (**Fig. 9C–F**). However, elastic modulus was strongly correlated with callus volume (*R* = 0.87, *p* < 0.0001), indicating that aberrant callus expansion in high-strain groups does not effectively restore structural integrity.

**Figure 9.**
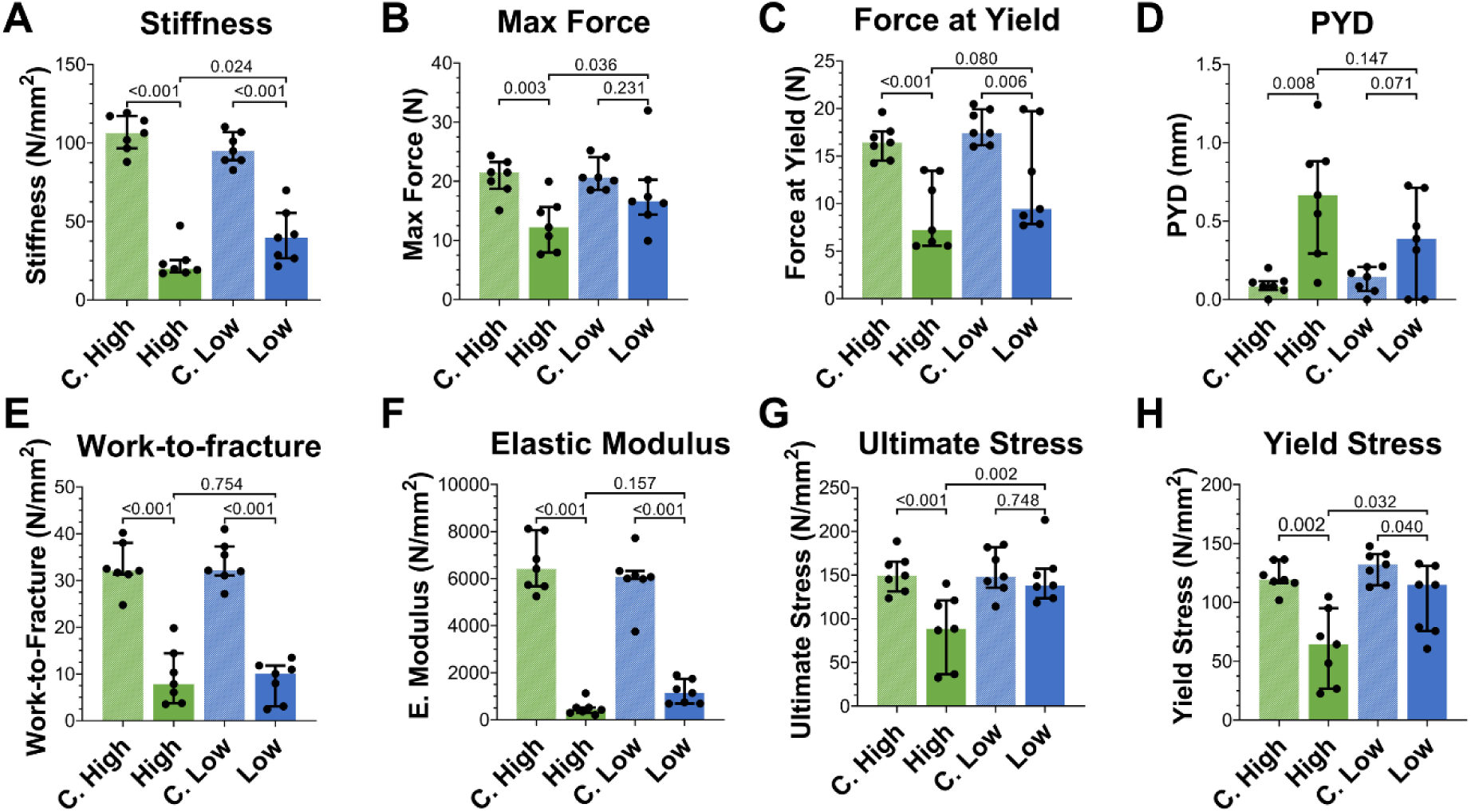
Interfragmentary Strain Determines Biomechanical Outcomes of Fracture Healing. Three-point bending analysis of femurs at 3 weeks post-fracture. **(A–D)** Whole-bone properties: stiffness (A), maximum force (B), force at yield (C), and post-yield displacement (PYD, D). **(E–H)** Tissue-level properties: work-to-fracture (E), elastic modulus (F), ultimate stress (G), and yield stress (H). Low-strain (high stiffness nail, <5% interfragmentary strain) groups significantly outperformed high-strain (low-stiffness nail, >15% strain) groups in stiffness (p = 0.024), maximum force (p = 0.036), ultimate stress (p = 0.002), and yield stress (p = 0.032). Data are median ± interquartile range; n = 7 per group. Contralateral controls (C. High, C. Low) represent uninjured femurs.

**Table 2.** Assessment of fracture healing outcomes. Comparative evaluation of whole-bone and tissue-level mechanical properties of femurs subjected to 3-point bending testing. Compared groups are high-strain (low-stiffness nail), low-strain (high-stiffness nail), control high (contralateral femur from high-strain group), and control low (contralateral femur from low-strain group). Values are presented as mean±SD. n = 7 per group.

|  | **Control High** | **High-Strain** | **Control Low** | **Low-Strain** | **High vs. Low p-value** | **C. High vs. High p-value** | **C. Low vs. Low p-value** |
| --- | --- | --- | --- | --- | --- | --- | --- |
| **Stiffness (N/mm)** | 106.16 ± 11.5 | 24.17 ± 10.7 | 96.26 ± 10.2 | 40.48 ± 17.2 | 0.024* | <0.001* | <0.001* |
| **Max load (N)** | 20.64 ± 3.1 | 12.72 ± 4.4 | 21.11 ± 2.6 | 18.19 ± 6.9 | 0.036* | 0.003* | 0.231 |
| **Force at Yield (N)** | 16.49 ± 1.8 | 8.95 ± 3.7 | 18.08 ± 1.8 | 12.39 ± 5.4 | 0.080 | <0.001* | 0.006* |
| **Post-yield displace. (mm)** | 0.09 ± 0.1 | 0.66 ± 0.4 | 0.12 ± 0.1 | 0.38 ± 0.3 | 0.147 | 0.008* | 0.071 |
| **Work-to-fracture (Nmm)** | 32.94 ± 5.02 | 9.39 ± 6.0 | 33.72 ± 4.6 | 8.54 ± 4.3 | 0.754 | <0.001* | <0.001* |
| **Elastic Modulus (N/mm2)** | 6582.70 ± 1150.7 | 490.80 ± 303.1 | 5999.99 ± 1166.7 | 1170.71 ± 503.2 | 0.157 | <0.001* | <0.001* |
| **Ultimate Stress (N/mm2)** | 152.07 ± 22.01 | 88.60 ± 41.4 | 153.46 ± 25.8 | 148.06 ± 31.7 | 0.002* | <0.001* | 0.748 |
| **Yield Stress (N/mm2)** | 121.60 ± 12.1 | 61.89 ± 31.7 | 130.64 ± 13.3 | 101.27 ± 29.0 | 0.032* | 0.002* | 0.040* |

### 5. Mechanically Unstable Fibrotic Healing Is Associated with Subtle Systemic Immune Changes

To evaluate whether mechanically impaired healing is accompanied by systemic immune alterations, peripheral blood immune populations were longitudinally profiled following fracture (**Fig. 10**). Within one hour post-fracture, both groups showed declines in circulating B cells and monocytes/macrophages and increases in total T cells and neutrophils, indicating a system-wide acute inflammatory response. By day 1, the high-strain group exhibited a 9.4% increase in B cells (72.63±4.72% vs. 63.27±9.43%, *p* = 0.001, **Fig. 10A(ii)**) and a 6.8% decrease in total CD3+ T cells (19.14±5.59% vs. 25.97±8.03%, p = 0.004, **Fig. 10A(iii)**) compared to the low-strain group. Among T cell subsets, CD4+ T helper cells were the only population with significant differences between groups, with a modest 2.6% increase in the high-strain group at hour 1 (19.53±4.83% vs. 16.7±2.33%, *p* = 0.032, **Fig. 10A(iv)**) and a 3.0% decrease at day 1 (9.99±2.46% vs. 12.96±3.15%, *p* = 0.025, **Fig. 10A(iv)**) compared to the low-strain group. These early differences attenuated over time, and no sustained between-group differences were observed at later time points. At day 21, callus tissue immunophenotyping revealed a lower myeloid proportion in the high-strain compared to the low-strain group (38.57±11.55% vs. 52.9±12.12%, *p* = 0.043), whereas the distributions of other immune cell subtypes were similar between groups (**Table 3**).

**Figure 10.**
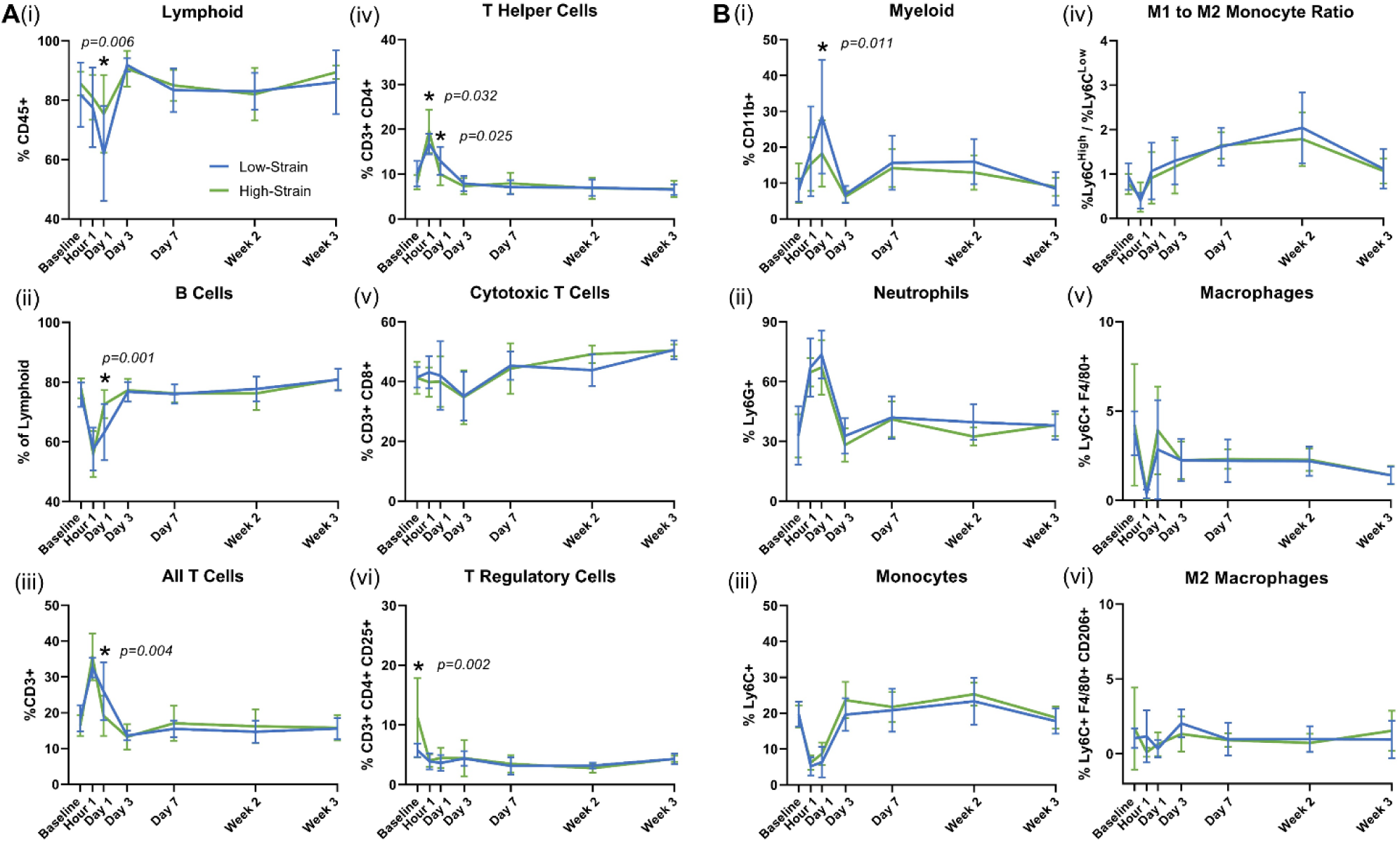
Interfragmentary Strain Shifts Systemic Immune Profile Post-Fracture. (**A**) Lymphoid cell dynamics: (**i**) Total CD45+ lymphoid cells, (**ii**) B cells, (**iii**) CD3+ T cells, (**iv**) CD4+ T helper cells, (**v**) CD8+ cytotoxic T cells, and (**vi**) CD25+ Tregs in peripheral blood. **(B)** Myeloid cell dynamics: (**i**) Total CD11+ myeloid cells, (**ii**) Ly6G+ neutrophils, (**iii**) Ly6C+ classical monocytes, (**iv**) M1 to M2 monocyte ratio, (**v**) F4/80+ macrophages, and (**vi**) CD206+ M2 macrophages. Low-strain groups (high-stiffness nail, <5% interfragmentary strain, blue) exhibit early CD4+ T cell expansion and B cell contraction, while high-strain groups (low-stiffness nail, >15% interfragmentary strain, green) maintain innate dominance. Data are mean ± SEM; n = 7 per group; Baseline values were captured 5 days before fracture.

**Table 3.**
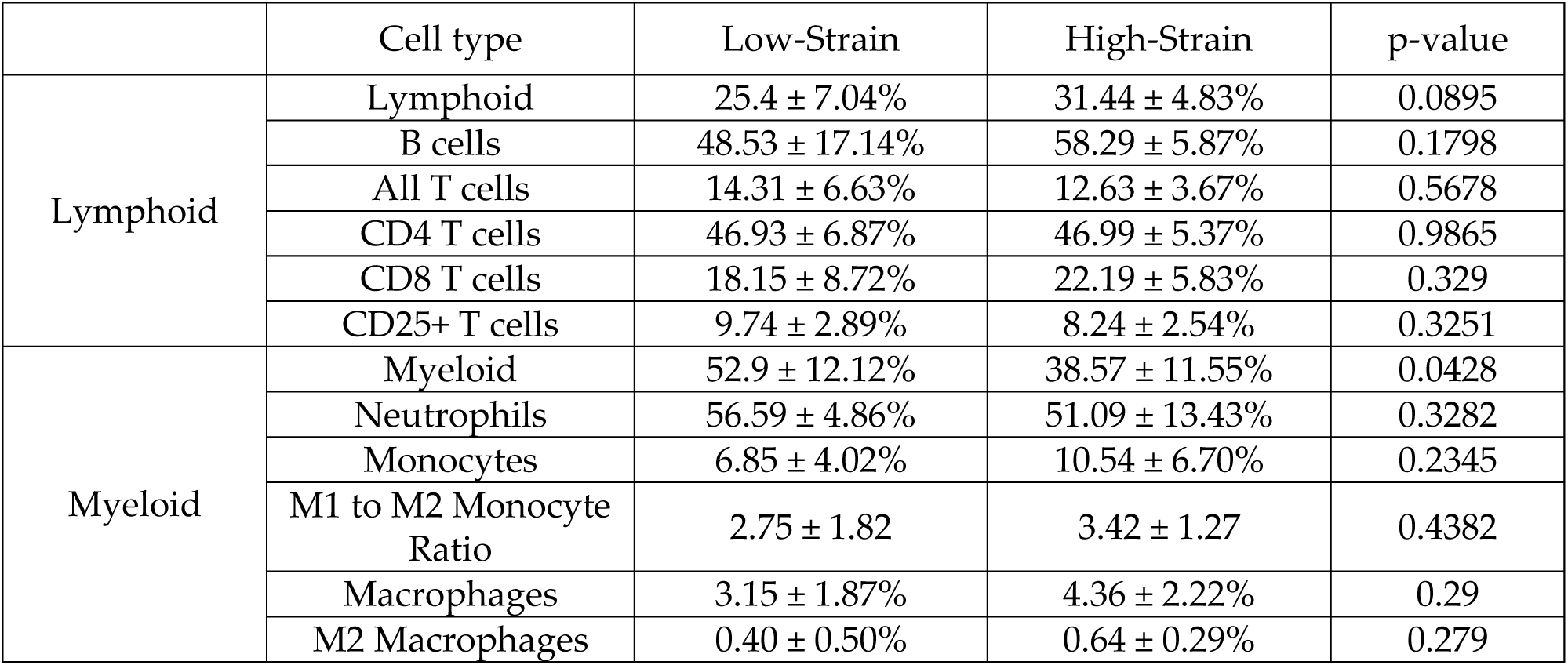
Immune cell population percentages in day-21 fracture callus. Comparative evaluation of callus population percentages between low-strain (high-stiffness nail) and high-strain (low-stiffness nail) fractures using flow cytometry. Values are presented as mean±SD. n = 7 per group.

|  | Cell type | Low-Strain | High-Strain | p-value |
| --- | --- | --- | --- | --- |
| Lymphoid | Lymphoid | 25.4 ± 7.04% | 31.44 ± 4.83% | 0.0895 |
|  | B cells | 48.53 ± 17.14% | 58.29 ± 5.87% | 0.1798 |
|  | All T cells | 14.31 ± 6.63% | 12.63 ± 3.67% | 0.5678 |
|  | CD4 T cells | 46.93 ± 6.87% | 46.99 ± 5.37% | 0.9865 |
|  | CD8 T cells | 18.15 ± 8.72% | 22.19 ± 5.83% | 0.329 |
|  | CD25+ T cells | 9.74 ± 2.89% | 8.24 ± 2.54% | 0.3251 |
| Myeloid | Myeloid | 52.9 ± 12.12% | 38.57 ± 11.55% | 0.0428 |
|  | Neutrophils | 56.59 ± 4.86% | 51.09 ± 13.43% | 0.3282 |
|  | Monocytes | 6.85 ± 4.02% | 10.54 ± 6.70% | 0.2345 |
|  | M1 to M2 Monocyte Ratio | 2.75 ± 1.82 | 3.42 ± 1.27 | 0.4382 |
|  | Macrophages | 3.15 ± 1.87% | 4.36 ± 2.22% | 0.29 |
|  | M2 Macrophages | 0.40 ± 0.50% | 0.64 ± 0.29% | 0.279 |

Additionally, the plasma fraction was profiled using a 32-analyte Milliplex MAP Mouse Cytokine/Chemokine Panel across the 1-hour to day 7 timepoints. Consistent with the peripheral blood immunophenotyping, cytokine and chemokine levels exhibited only subtle differences, with no significant between-group changes detected (**Supplementary Table S3**). While these findings are subtle, specimen-level and multivariate analyses help delineate the differences and identify several potential biomarkers for fracture healing quality.

### 6. Systemic and Local Immune Signatures Predict Biomechanical Healing Outcomes

Specimen-level linear regression identified 34 immune factors, including cytokines, chemokines, and cell populations, that were associated with functional healing outcomes (**Table 4**). Early after injury, MIG (CXCL9) levels showed an inverse association with maximum force at 1 hour in peripheral blood (R = −0.656, *p = 0.051*) and a stronger inverse association at day 1 (R = −0.786, p = 0.003; **Fig. S3**). At day 7 in the peripheral blood, MIP-1α (Macrophage Inflammatory Protein-1α) correlated positively with elastic modulus (R = 0.692, *p = 0.020*), and M-CSF (Macrophage Colony-Stimulating Factor) correlated positively with bending stiffness (R = 0.607, *p = 0.013*) (**Fig. S3**).

**Table 4.**
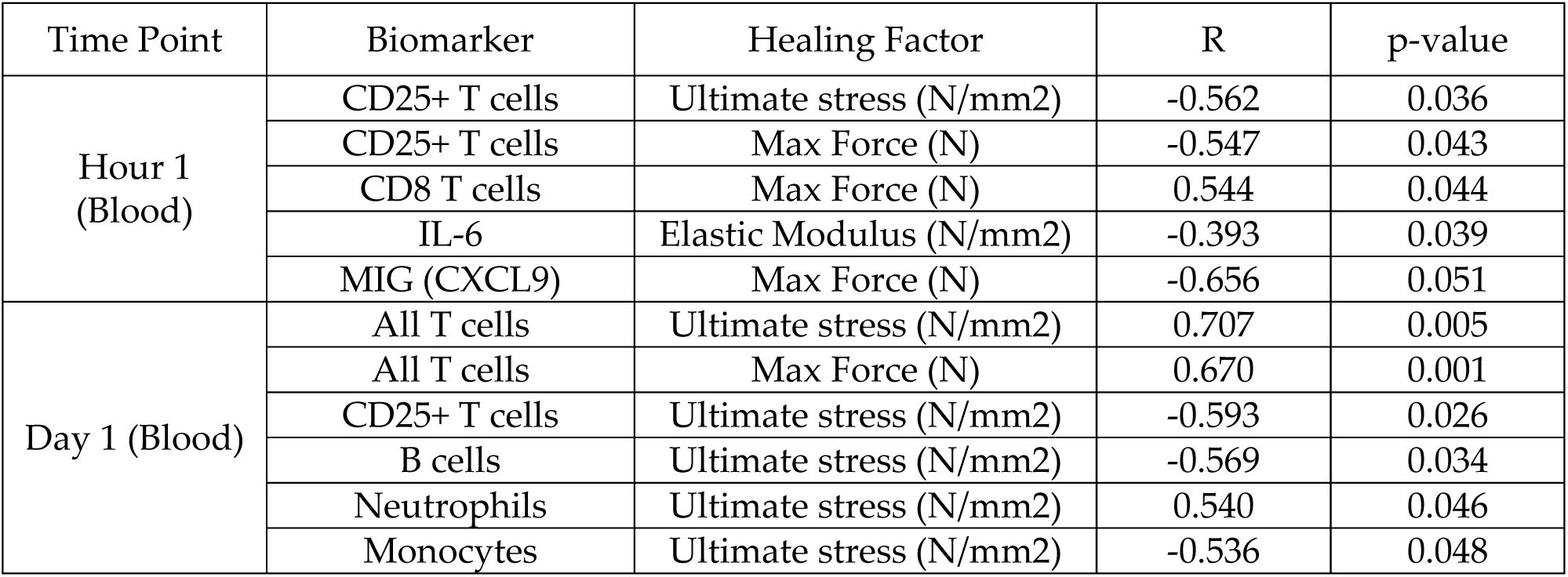
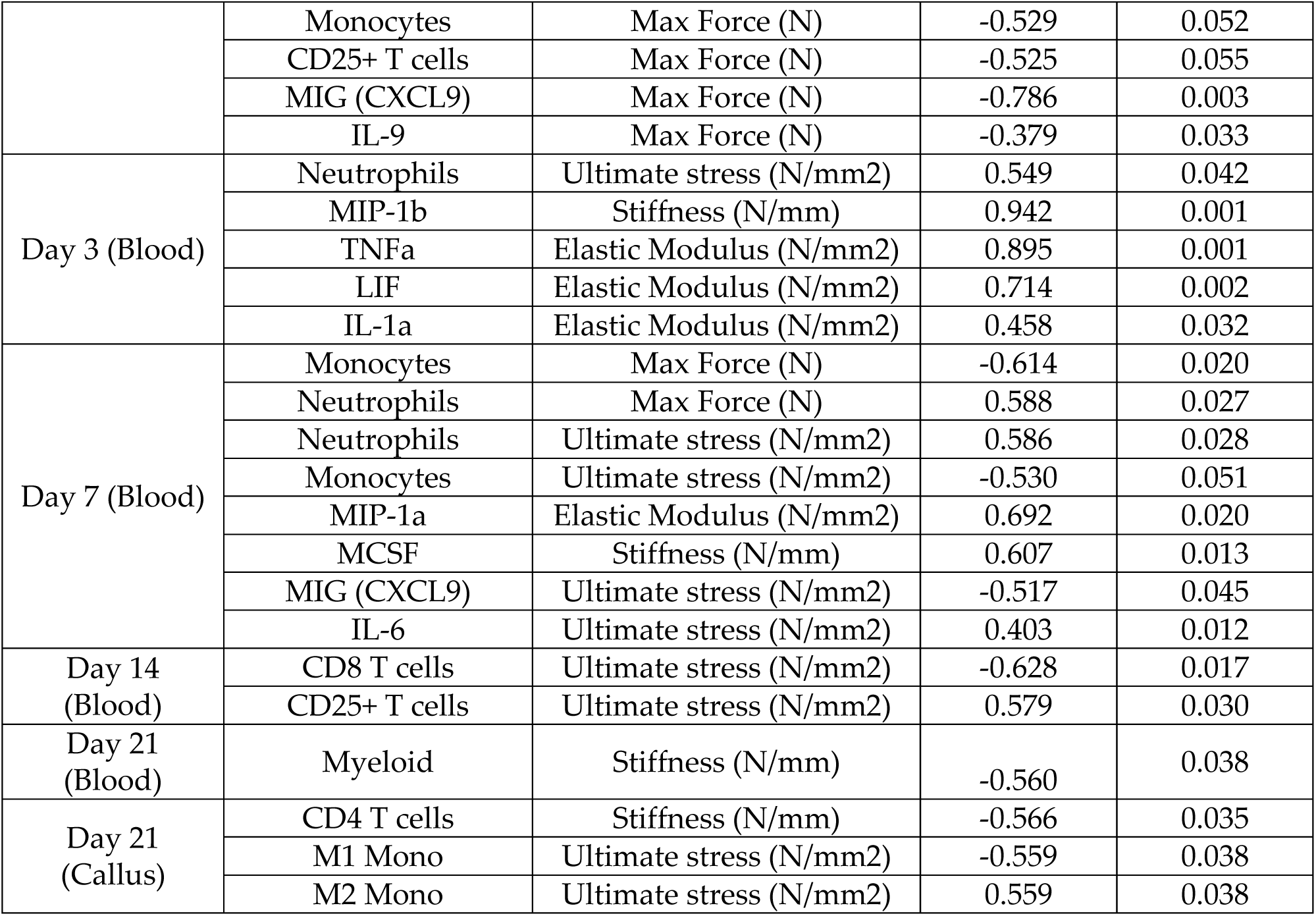
Correlations between immune factors and healing outcomes.

| Time Point | Biomarker | Healing Factor | R | p-value |
| --- | --- | --- | --- | --- |
| Hour 1 (Blood) | CD25+ T cells | Ultimate stress (N/mm2) | -0.562 | 0.036 |
|  | CD25+ T cells | Max Force (N) | -0.547 | 0.043 |
|  | CD8 T cells | Max Force (N) | 0.544 | 0.044 |
|  | IL-6 | Elastic Modulus (N/mm2) | -0.393 | 0.039 |
|  | MIG (CXCL9) | Max Force (N) | -0.656 | 0.051 |
| Day 1 (Blood) | All T cells | Ultimate stress (N/mm2) | 0.707 | 0.005 |
|  | All T cells | Max Force (N) | 0.670 | 0.001 |
|  | CD25+ T cells | Ultimate stress (N/mm2) | -0.593 | 0.026 |
|  | B cells | Ultimate stress (N/mm2) | -0.569 | 0.034 |
|  | Neutrophils | Ultimate stress (N/mm2) | 0.540 | 0.046 |
|  | Monocytes | Ultimate stress (N/mm2) | -0.536 | 0.048 |
|  | Monocytes | Max Force (N) | -0.529 | 0.052 |
|  | CD25+ T cells | Max Force (N) | -0.525 | 0.055 |
|  | MIG (CXCL9) | Max Force (N) | -0.786 | 0.003 |
|  | IL-9 | Max Force (N) | -0.379 | 0.033 |
| Day 3 (Blood) | Neutrophils | Ultimate stress (N/mm2) | 0.549 | 0.042 |
|  | MIP-1b | Stiffness (N/mm) | 0.942 | 0.001 |
|  | TNFa | Elastic Modulus (N/mm2) | 0.895 | 0.001 |
|  | LIF | Elastic Modulus (N/mm2) | 0.714 | 0.002 |
|  | IL-1a | Elastic Modulus (N/mm2) | 0.458 | 0.032 |
| Day 7 (Blood) | Monocytes | Max Force (N) | -0.614 | 0.020 |
|  | Neutrophils | Max Force (N) | 0.588 | 0.027 |
|  | Neutrophils | Ultimate stress (N/mm2) | 0.586 | 0.028 |
|  | Monocytes | Ultimate stress (N/mm2) | -0.530 | 0.051 |
|  | MIP-1a | Elastic Modulus (N/mm2) | 0.692 | 0.020 |
|  | MCSF | Stiffness (N/mm) | 0.607 | 0.013 |
|  | MIG (CXCL9) | Ultimate stress (N/mm2) | -0.517 | 0.045 |
|  | IL-6 | Ultimate stress (N/mm2) | 0.403 | 0.012 |
| Day 14 (Blood) | CD8 T cells | Ultimate stress (N/mm2) | -0.628 | 0.017 |
|  | CD25+ T cells | Ultimate stress (N/mm2) | 0.579 | 0.030 |
| Day 21 (Blood) | Myeloid | Stiffness (N/mm) | -0.560 | 0.038 |
| Day 21 (Callus) | CD4 T cells | Stiffness (N/mm) | -0.566 | 0.035 |
|  | M1 Mono | Ultimate stress (N/mm2) | -0.559 | 0.038 |
|  | M2 Mono | Ultimate stress (N/mm2) | 0.559 | 0.038 |

Within innate immune compartments, peripheral blood neutrophil proportions at day 7 were positively correlated with maximum force (R = 0.59, *p = 0.027*) and ultimate stress (R = 0.59, *p = 0.028*). In callus tissue at day 21, monocyte polarization states exhibited opposing associations with mechanical performance: M1-like monocytes were negatively correlated with ultimate stress (R = −0.56, *p = 0.037*), whereas M2-like monocytes were positively correlated (R = 0.59, *p = 0.037*).

Temporal patterns in adaptive immune populations further revealed stage-specific associations with healing outcomes (**Fig. 11**). At day 1, circulating CD3⁺ T cells were positively associated with ultimate stress (R = 0.71, *p = 0.005*; **Fig. 11A(i)**), whereas CD25⁺ regulatory T cells showed a negative association (R = −0.59, *p = 0.026*; **Fig. 11A(ii)**). By day 14, circulating CD8⁺ T cells were negatively correlated with ultimate stress (R = −0.62, *p = 0.017*; **Fig. 11B(i)**), while regulatory T cells (Tregs) demonstrated a positive association (R = 0.58, *p = 0.030*; **Fig. 11B(ii)**). In callus tissue at day 21, CD4⁺ T cell abundance was inversely associated with stiffness (R = −0.57, p = 0.034).

**Figure 11.**
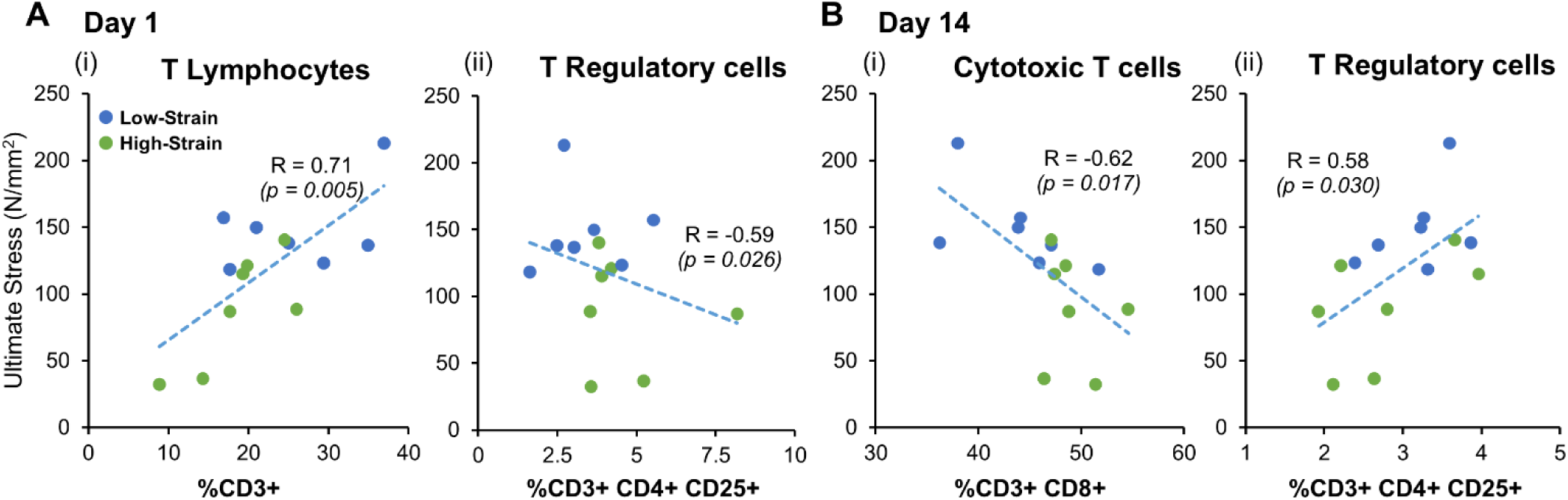
Systemic Immune Profile Predicts Biomechanical Healing Outcomes. **(A)** Day 1 post-fracture: (**i**) Positive correlation between CD3+ T cell frequency and ultimate stress (R = 0.71, p = 0.005). (**ii**) Negative correlation between CD25+ Treg frequency and ultimate stress (R = -0.59, p = 0.026). (**B**) Day 14 post-fracture: (**i**) Negative correlation between CD8+ T cell frequency and ultimate stress (R = -0.62, p = 0.017). (**ii**) Positive correlation between CD25+ Treg frequency and ultimate stress (R = 0.58, p = 0.030). Data are linear regression fits, n = 7 per group. Green dots represent individual high-strain (low-stiffness nail, >15% interfragmentary strain) samples, blue dots represent individual low-strain (high-stiffness nail, <5% interfragmentary strain) samples.

Collectively, these findings identify temporally structured relationships among soluble mediators, innate immune cells, adaptive immune populations, and biomechanical measures of fracture repair. These associations support the potential utility of integrated immune profiling for identifying candidate biomarkers of fracture healing quality.

### 7. Circulating Immune Phenotypes Predict Functional Outcomes Under Mechanical Instability

To identify multivariate immune features associated with repair quality, partial least squares regression (PLSR) was applied to longitudinal peripheral blood immune profiles (1 hour to 21 days post-fracture) in relation to biomechanical outcomes. This approach revealed distinct immune trajectories that differentiated high-strain and low-strain conditions, with peak model performance observed at 1 hour (R = 0.71, *p = 0.005*), day 3 (R = 0.57, *p = 0.032*), and day 14 (R = 0.72, *p = 0.004*) post-injury (**Fig. 12**).

**Figure 12.**
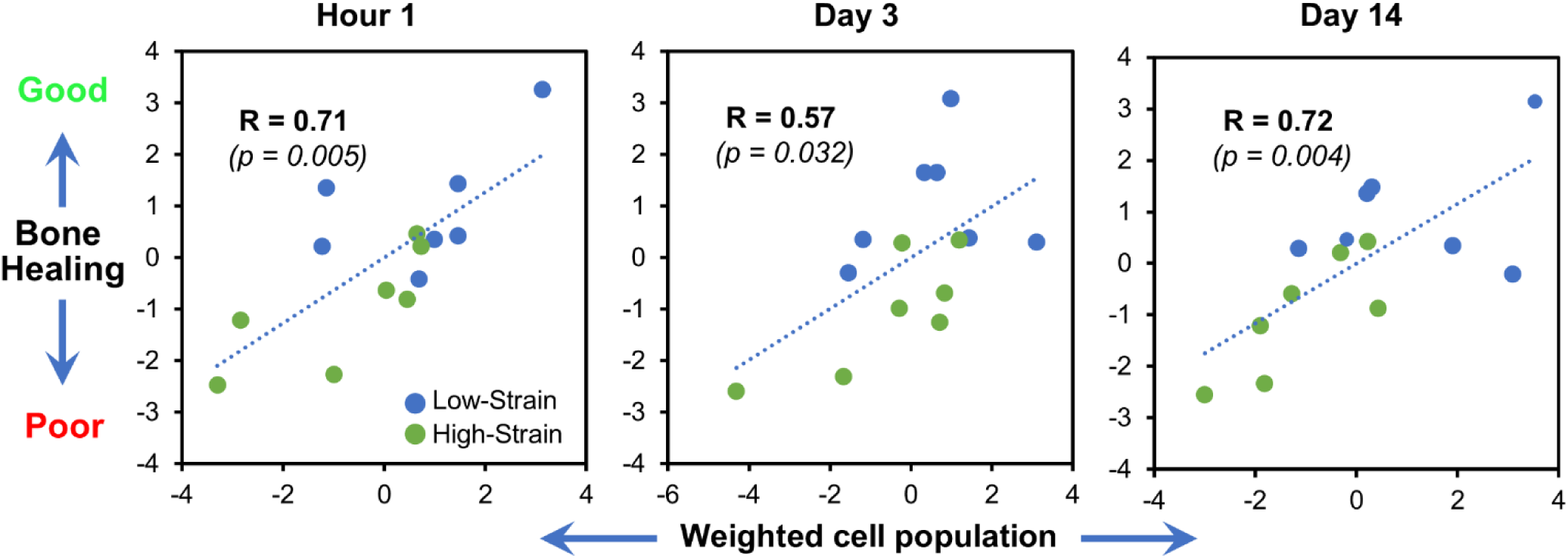
Early Immune Profiles Predict Biomechanical Outcomes of Fracture Healing. Partial least squares regression (PLSR) analysis of systemic immune cell populations (X matrix) against functional healing outcomes (Y matrix: stiffness, maximum force, ultimate stress, elastic modulus). PLSR score plot demonstrates maximal separation between high-strain (low-stiffness nail, >15% interfragmentary strain, green) and low-strain (high-stiffness nail, <5% interfragmentary strain, blue) groups at 1 hour, day 3, and day 14 post-fracture. Weighted coefficient values in the model identify CD206+ macrophages (1 hour, day 14) and CD25+ Tregs (day 14) as top predictors. n = 7 per group.

Model loadings identified temporally structured predictors of healing outcomes. At early time points, CD206⁺ macrophages at 1 hour (weighted coefficient = 0.228) and F4/80⁺ macrophages at day 3 (weighted coefficient = 0.206) were the most influential contributors to model predictions. At later stages, CD25⁺ regulatory T cells at day 14 emerged as the dominant predictor (weighted coefficient = 0.958), alongside CD206⁺ macrophages (weighted coefficient = 0.233), indicating a shift in the relative contribution of immune subsets over time.

These results demonstrate that circulating immune profiles contain temporally resolved information associated with biomechanical healing outcomes. The observed transition from early innate immune signals to later adaptive immune features is consistent with a staged immunological response that may reflect the progression of fracture repair under differing mechanical environments.

## C. Discussion

### 1. A Murine Model of Strain-Dependent Hypertrophic Nonunion

Hypertrophic nonunion, commonly attributed to mechanical instability, remains difficult to diagnose early and lacks effective non-surgical treatment strategies. Existing preclinical models predominantly represent atrophic nonunion or segmental defects and therefore do not capture the strain-dependent pathophysiology observed in clinical hypertrophic nonunion (*34*). In this study, a murine model was developed to impose controlled interfragmentary strain, enabling direct investigation of how mechanical instability influences fracture repair.

Interfragmentary strain is a key determinant of fracture stability and regulates multiple aspects of healing, including inflammation, vascularization, and tissue differentiation (*35*). Excessive strain disrupts these processes and has been associated with fibrotic repair and nonunion (*14*). Using superelastic nitinol (*36, 37*), intramedullary implants with defined stiffness profiles were engineered to generate distinct mechanical environments. Low-stiffness fixation produced high strain (15–30%), whereas high-stiffness fixation limited strain to <5%, approximating clinically stable conditions.

High-strain fractures (low-stiffness nail, >15% strain) formed larger calluses but exhibited reduced biomechanical integrity compared to low-strain controls (high-stiffness nail, <5% strain). This dissociation between callus size and functional strength is consistent with clinical observations in hypertrophic nonunion, where tissue accumulation does not translate to effective load-bearing (*38*). Importantly, increased bone volume (BV) in the high-strain group occurred alongside proportional increases in total callus volume (TV), resulting in unchanged bone volume fraction (BV/TV). Similarly, bone mineral density (BMD) did not differ between groups. These findings indicate that impaired healing arises from altered spatial organization and tissue composition rather than reduced mineralization capacity. This model, therefore, isolates mechanical instability as a primary driver of maladaptive repair and provides an experimentally controlled system for studying strain-dependent mechanisms.

### 2. Mechano-Immune Regulation of Fibrotic Remodeling

Mechanical instability alters the fracture microenvironment in ways that extend beyond structural effects to include immune and stromal interactions (*39–41*). Spatial transcriptomic analyses identified fibroblast-enriched regions within high-strain calluses that co-localized with macrophages expressing *Lgals3* and osteoclasts expressing *Mmp9*. These cellular niches were characterized by elevated expression of genes associated with fibrosis, including *Spp1*, *Pdgfrb,* and *Lgals3*, suggesting coordinated macrophage-fibroblast signaling. Macrophages expressing *Lgals3* have been implicated in chronic fibrotic responses in other tissues, supporting the relevance of this phenotype (*42*).

Computational inference of cell–cell communication implicated a thrombospondin-syndecan-4 signaling axis, in which macrophage-derived thrombospondins (*Thbs1/2*) interact with fibroblast syndecan-4 receptors. Syndecan-4 is a mechanosensitive co-receptor involved in focal adhesion dynamics and extracellular matrix organization (*43*). Increased deposition of thrombospondin-1 in high-strain calluses, as confirmed by immunohistochemistry, supports activation of this pathway *in vivo*. These findings are consistent with prior studies demonstrating roles for thrombospondins in fibrotic remodeling across tissues (*44, 45*).

In parallel, osteoclast-associated *Mmp9* expression may contribute to activation of latent TGF-β, further promoting fibroblast activation and matrix deposition (*46, 47*). Gene ontology analyses supported enrichment of pathways related to extracellular matrix organization and fibroblast regulation, with macrophages as prominent contributors. Together, these observations support a model in which mechanical instability sustains a feedback loop involving macrophages, fibroblasts, and matrix remodeling processes, resulting in persistence of fibrotic tissue and impaired ossification.

### 3. Systemic Immune Correlates of Strain-Dependent Healing

In addition to local tissue changes, fracture healing is associated with systemic immune responses that may reflect or influence repair outcomes. Prior human studies have reported associations between circulating immune profiles and delayed healing, including enrichment of CD8⁺ effector populations (*48*). Consistent with these observations, the present study identified persistent CD8⁺ T cell signatures and reduced representation of regulatory T cells (CD25⁺ Tregs) under high-strain conditions, suggesting a systemic inflammatory state associated with impaired repair.

Comparative findings from murine aging models further support the association between sustained inflammation and delayed healing, including enrichment of CD8⁺ cells and macrophages in poorly healing calluses (*49*). These convergent observations suggest that distinct immune profiles may characterize suboptimal healing trajectories across different biological contexts.

Multivariate analysis using partial least squares regression identified temporally structured immune predictors of biomechanical outcomes. Early macrophage-associated signals, including CD206⁺ populations, and later adaptive immune features, particularly regulatory T cells, were among the strongest contributors to model performance. These findings indicate that fracture healing is associated with coordinated temporal changes in immune composition and that deviations from this pattern may be associated with impaired healing outcomes.

Collectively, these results suggest that circulating immune features reflect underlying strain-dependent repair processes and may provide a basis for developing prognostic biomarkers. The convergence of local fibrotic signatures and systemic immune profiles supports a link among the mechanical environment, immune regulation, and healing outcomes.

### 4. Limitations and Clinical Translation

Although this model reproduces key features of strain-driven hypertrophic nonunion, several limitations should be considered. First, murine fractures typically achieve union within 6–9 weeks, and thus the model represents delayed healing rather than permanent nonunion. This distinction is relevant because human nonunions frequently require surgical intervention and may not resolve spontaneously. However, the model captures an early pathogenic window during which mechanical instability promotes fibrosis and immune dysregulation, providing insight into processes that may precede chronic nonunion. In addition, biomechanical assessments at day 21 primarily reflect early hard-callus properties, whereas further increases in strength are expected with ongoing remodeling.

Second, clinical fracture healing occurs across a spectrum of mechanical environments influenced by implant selection, surgical technique, and patient-specific factors. The present model adopts a reductionist approach by varying fixation stiffness to isolate interfragmentary strain as a primary variable. While this enables controlled mechanistic investigation, it does not encompass the full heterogeneity of clinical fracture repair.

Third, interfragmentary strain was not continuously measured *in vivo*, limiting the ability to resolve dynamic relationships between mechanical loading and biological responses. Nevertheless, the observed outcomes are consistent with computational models predicting that elevated strain promotes fibrotic repair and impaired healing (*50*). Integration of strain-sensing technologies in future studies may improve the temporal resolution of mechanobiological interactions.

Fourth, although specific fibrotic pathways (THBS–SDC4, *Lgals3*) and immune populations (CD206⁺ macrophages, regulatory T cells) were identified, the pleiotropic nature of immune signaling suggests that additional mediators are likely involved. Expanded profiling of cytokine networks and stromal cell subsets may further refine the mechanistic framework.

Fifth, the exclusive use of male mice reduces variability related to hormonal cycling but limits generalizability. Sex-specific differences in immune function and bone biology warrant inclusion of female cohorts in future work.

Sixth, interspecies differences in immune responses and skeletal biology necessitate validation in larger animal models to assess translational relevance. In particular, confirmation of the THBS–SDC4 axis and associated immune biomarkers in systems with humanized immune components will be important for evaluating clinical applicability.

Finally, spatial transcriptomic analyses were conducted on a limited number of samples, which constrains generalizability. This limitation reflects current technical and resource constraints in spatial profiling. It was partially addressed through complementary histological and immunohistochemical validation on adjacent sections and by focusing interpretation on within-sample spatial relationships. Population-level inferences from these data should therefore be interpreted with caution.

### 5. Summary and Conclusion

This study establishes a murine model in which controlled mechanical instability drives delayed fracture healing characterized by fibrotic remodeling and reduced biomechanical integrity. By isolating interfragmentary strain as a primary determinant, the model enables direct investigation of how mechanical cues shape immune–stromal interactions during repair.

The findings support a mechanistic association between elevated strain, macrophage–fibroblast signaling, and persistence of fibrotic tissue, with evidence implicating THBS–SDC4 signaling and *Lgals3*-associated macrophage activity. Consistent with this local fibrotic phenotype, circulating immune features, including CD206⁺ macrophages and CD25⁺regulatory T cells, were associated with biomechanical outcomes, suggesting their potential utility as prognostic indicators.

Together, these results provide a framework that links the mechanical environment, immune regulation, and healing outcome. This integrative perspective may inform future strategies for early risk stratification and the development of interventions targeting strain-dependent pathways in fracture repair.

## D. Methods

### 1. Animal model and surgical procedures

All animal procedures adhered to the guidelines approved by the Institutional Animal Care and Use Committee (IACUC) at the University of Kentucky. Adult male C57BL/6 mice (22–24 weeks old, The Jackson Laboratory) were used in this study. Prior to any procedures, animals underwent a minimum 7-day acclimatization period. The mice were randomly assigned to two experimental groups receiving either low- or high-stiffness intramedullary nails (n = 11 per group). The intramedullary nails were custom manufactured from nitinol rods (Chamfr), with diameters ranging from 0.31 to 0.51 mm to modulate stiffness.

An established transverse fracture model (*32*) was employed to create femoral fractures. This method does not require any incisions through muscle and mimics fractures resulting from blunt trauma in humans with reproducible precision (*51, 52*). Briefly, mice were anesthetized (1.5 mg/kg ketamine + 1.5 mg/kg xylazine, based on body mass of mice), a bland ophthalmic ointment was applied, and the surgical site was shaved and wiped with 70% ethanol, followed by 10% povidone-iodine solution. A small incision was made at the patella and laterally dislocated to expose the knee joint. Using a 0.5 mm centering bit, a hole was made in the intercondylar notch and reamed with either a 25- or 27-gauge hypodermic needle to ensure precise fixation of the low- and high-stiffness nails. An intramedullary guide tube (NiTi, 0.2 mm diameter) was then inserted into the femur. With the guide tube in place, the right leg was positioned under the guillotine (blunt striker) of the murine fracture apparatus (RISystems). The blunt striker was then released to produce impact kinetic energy (*E_k_*) of ∼40 *mJ*, which has been shown to create a transverse fracture in adult C57BL/6 mice (*53*). The guide tube was then replaced with the appropriate NiTi intramedullary nails to stabilize the femur. The skin at the patella was sutured with 6-0 vicryl suture (Ethicon). Then, 1 mL of saline was administered intraperitoneally for hydration, followed by subcutaneous administration of buprenorphine SR (1.5 mg/kg body mass) for analgesia. The animals were then placed into a recovery cage and monitored as per the established post-operative animal care protocols. Animals were monitored daily for 7 days and weekly thereafter to ensure post-operative recovery. Day 21 was chosen as the primary end point to coincide with the hard-callus/early remodeling stage in mice, when mineralized bridging is evident but remodeling is incomplete, enabling detection of strain-dependent differences in tissue quality and mechanics while preserving feasibility for mechanistic assays (3–5-week “late” window in mice) (*33*). To minimize variance from sex-dependent differences in fracture repair and hormone-responsive immune/osteogenic programs, only male mice were used in this study. Each experimental group comprised 11 mice. Of these, 7 mice were allocated for destructive three-point bending tests; the same 7 were also used for longitudinal peripheral blood flow cytometry and multiplex analyses, and their callus tissue was collected at day 21 for flow cytometry after three-point bending. The remaining 4 mice were dedicated to histological analyses. All 11 mice per group were evaluated by micro-CT.

### 2. Tissue collection and processing

Blood samples were collected longitudinally from mice in K2 EDTA Microvette tubes (Sarstedt) from the submental and submandibular veins, following an alternating schedule. Baseline samples were obtained five days prior to surgery, with subsequent samples taken at 1 hour, 1, 3-, 7-, 14-, and 21-days post-surgery. Blood samples were centrifuged at 900 g for 5 minutes to separate cells from plasma. The supernatant containing plasma was stored at -80°C until analyzed using a multiplex assay kit. The cellular fraction of the blood was then processed, stained, and analyzed using flow cytometry.

For analyzing callus tissue cell populations, callus tissue at day 21 was minced, digested in a 0.1% collagenase solution (MP Millitec) at 37°C for 2 hours, neutralized with Dulbecco’s modified Eagle’s media (DMEM, Gibco) containing 10% fetal bovine serum (FBS, Gibco), filtered through a 100 µm cell strainer, and centrifuged at 400 g for 5 minutes to remove the supernatant. The cell pellet was then processed, stained, and analyzed using flow cytometry.

At 21 days post-surgery, the animals were euthanized, and their femurs (fractured and contralateral control) were collected in DMEM. The bones were subjected to morphometric analysis and biomechanical testing at room temperature. For histological analyses, the samples were then fixed in 10% neutral buffered formalin (ThermoFisher Scientific) for 24 hours, stored in 10 mM PBS at 4°C, and analyzed as described below.

### 3. MicroCT and morphometric analysis

Fracture healing outcomes were assessed at week 3 post-surgery using microcomputed tomography (microCT), as described (*54*). To prevent any interference from the intramedullary nails, they were meticulously removed before the scanning process. A 360° scan was performed using a high-resolution small animal imaging system, SkyScan1276 microCT (Bruker, Billerica, MA), with a 1 mm aluminum filter, with the voltage set to 70 kV and image pixel size 20.32 µm. Images were then reconstructed (NRecon, Bruker) with smoothing (set to 3), ring artifact correction (set to 4), and beam hardening correction (set to 30%). In preparation for morphometric analysis, cylindrical phantoms with diameters of 2 mm and known bone mineral densities of 0.25 and 0.75 g/cm³ were submerged in deionized water, scanned, and reconstructed using the same settings.

The cortical region of the bone was delineated using a five-level Otsu algorithm. The threshold values for each level were determined based on the contralateral femur. Level 3 was scrutinized against the raw greyscale images to ensure the accuracy of the threshold, and it was subsequently employed as the cutoff threshold for analyzing the cortical region of the fractured bone. A 5 mm section (±2.5 mm from the fracture) at the fracture site was designated as the volume of interest (VOI), with the cortical bone being defined as the region of interest (ROI). The ROI was processed algorithmically within the CTan software by Bruker to ensure consistent and precise analysis of both the callus and cortical bone separately. An adaptive threshold (Mean-C) was applied to the callus region to accommodate variations in mineral density and ensure alignment with the raw greyscale images. Three-dimensional analyses were then carried out on the callus and cortical sections using these chosen thresholds. The CTvox software (Bruker) was utilized to construct three-dimensional volume-rendered images of the bone.

### 4. Mechanical testing

We used three-point bending to evaluate whole-bone mechanical competence, as described previously (*55*). This assay offered high reproducibility and controlled loading at the diaphyseal callus, and it permitted derivation of tissue-level properties via beam theory, facilitating linkage between mechanical behavior and callus architecture/fibrosis. The fracture and contralateral femurs were weighed, and their dimensions were determined through digital analysis of microCT scans. The three-point bending tests were executed using a Universal Testing Machine (Instron) at a strain rate of 30 mm/min, with an 8 mm span length, all performed at room temperature. Comprehensive whole-bone properties, encompassing parameters like stiffness, maximum force, work-to-fracture, and post-yield displacement (PDY), were deduced from the stress-strain curve. Furthermore, tissue-level parameters such as elastic modulus, yield stress, and ultimate stress were computed employing beam theory equations as listed below.

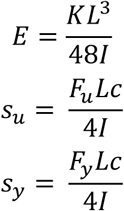

Where E = elastic modulus, K = stiffness, L = span length, I = moment of inertia, s_u_ = ultimate stress, F_u_ = ultimate force, c = distance from centroid of cross-section to outer most point on the cross-section, s_y_ = yield stress, and F_y_ = yield force. To calculate the average area moment of inertia (MOI) essential for these equations, a 5 mm segment (within ± 2.5 mm from the fracture site) was chosen from the microCT scans as the VOI, and a two-dimensional analysis was carried out.

The stiffness of the nitinol intramedullary nails was also measured through a three-point bend test performed at a strain rate of 10 mm/min with an 8 mm span length. Stiffness was calculated from the slope of the linear region. For both nail-only and nail-plus-femur tests, stiffness was determined directly from the force-displacement response, without requiring assumptions about bone material properties.

The flexure strain of the stabilized femur was utilized as a metric for measuring the interfragmentary movement. It was evaluated on fractured femora stabilized with the intramedullary nail under physiological loading conditions using three-point flexure testing. Because the mid-diaphyseal fracture renders the bone discontinuous, the nail is the principal load-bearing element. Therefore, strain was interpreted based on the nail’s cylindrical geometry rather than the surrounding bone tissue. Flexural strain was calculated using the standard expression for cylindrical beams:

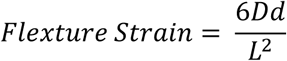

Where D = axial displacement, d = cylinder diameter, and L = span length.

A loading range of 1.4–2.1 N was selected to approximate physiological midshaft bending loads in the mouse femur, consistent with femur-specific load estimation reporting a bending moment of ∼3.5 ± 0.7 N·mm (*56*), which corresponds to ∼1.4–2.1 N in three-point bending with an 8 mm span. This range also agrees with hindlimb musculoskeletal modeling predicting joint and muscle resultants of similar magnitude (*57*). The strain measured within this force range was averaged over all tested samples.

### 5. Histology

For histological examination, mice were euthanized on day 21 post-surgery. Subsequently, the femurs with fractures were meticulously dissected and stripped of adjacent muscle tissues. The collected femurs then underwent decalcification using a solution containing 10% EDTA 4Na 2H_2_0 (Sigma ED4SS) and 9% EDTA 2Na 2H_2_0 (Sigma E5134) in sterilized deionized water. For samples designated for Visium spatial transcriptomics, 0.1% diethyl pyrocarbonate (Sigma D5758) was added to the decalcification solution to neutralize RNase enzymes. This decalcification solution was changed every 3 days over the course of 3 weeks. Following successful decalcification, the samples were paraffin-embedded and sectioned into 5 µm thick slices, which were subsequently mounted onto microscope slides (Apex Bond, Leica). The prepared slides were then subjected to staining using either a Movat’s pentachrome kit (StatLab) or a Picro-Sirius Red staining kit (StatLab), following the respective manufacturer’s instructions, or to immunohistochemistry (IHC) as described below.

Briefly, IHC was performed using the Ventana Discovery Ultra platform. 5 µm thick sections of formalin-fixed paraffin-embedded (FFPE) tissue were deparaffinized onboard, followed by on-board antigen retrieval as detailed below, primary antibody incubation at 37°C for 1 hour, detection as indicated below, and visualization with Ventana ChromoMap DAB.

SOX9 (Cell Signaling Technology 82630; 1:200) and SP7 (Thermo PA5-115697; 1:100) were retrieved using Ventana CC1 buffer under standard conditions. Primary antibody was amplified by incubation with Ventana anti-rabbit-HQ, then anti-HQ-HRP for 20 minutes each. CD-31 (Cell Signaling Technology 77699; 1:100) was retrieved using Ventana CC2 standard with primary amplification and detection as above. SMA (Abcam ab5694; 1:500) used CC1 standard and detection of primary antibody by Ventana OmniMap anti-rabbit-HRP. All slides were run concurrently with negative control and known positive control tissue to confirm assay performance.

TSP1 IHC was performed manually on 5 µm FFPE sections using an HRP/DAB kit (Abcam ab236469) without antigen retrieval to preserve tissue architecture. Sections were incubated with anti-Thbs1 (Invitrogen MA5-64220; 1:70). Briefly, endogenous peroxidase was blocked (10 min), protein block applied (30 min), primary antibody incubated overnight, HRP conjugate applied (30 min), DAB developed (15 min), then slides were counterstained, dehydrated, and coverslipped. All slides were run with negative control and known positive control tissue to confirm assay performance.

Stained slides were imaged on an Axioscan Z7 with pentachrome and IHC-stained slides being imaged under bright field light and picrosirius red slides being imaged under both bright field and polarized light. Immunohistochemical quantification was conducted using QuPath (v0.5.1). For each sample, 3–5 regions of interest (500 × 500 pixels) were manually placed within either the intramedullary or periosteal callus compartments, with selection prioritized along the fracture line within a ±2 mm margin. Positive cell percentages (positive cell detection module) or positive tissue area (DAB threshold) were calculated within each region and averaged per sample to account for tissue heterogeneity. Group-level means were then derived from 3–4 representative slides per condition, excluding any slides exhibiting significant tissue detachment. The fibrotic area was measured using Zeiss image analysis software. Only slides including the center section of the femur and fracture callus were used for analysis. The fibrotic tissue areas were manually selected with the area calculation tool and summed together to get the total fibrotic area per slide. Fibrotic areas were defined by the presence of fibrillar collagen visible under polarized light. Fibrotic areas were only measured if they were inside the fracture callus or within the fracture gap in the medullary canal. To quantify collagen architecture, high-magnification polarized images of each fibrotic area were analyzed in CT-FIRE v3 (MATLAB Runtime R2023b; default parameters) (*58*); images were converted to greyscale by CT-FIRE and ROIs were drawn to exclude bone and cartilage. The following fiber metrics were extracted and summarized per section and per animal for statistical analysis: width (mean width along the fiber centerline), angle (orientation relative to the positive x-axis, 0–180°), and length (end-to-end contour length).

### 6. Spatial Transcriptomics

We prepared libraries for 10X Genomics Visium CytAssist following the manufacturer’s guidelines. Formalin-fixed paraffin-embedded (FFPE) tissue sections (5 µm thick) were positioned on charged glass slides and baked at 42°C for 3 hours. After deparaffinization and hematoxylin and eosin (H&E) staining, high-resolution imaging was performed using a Leica Aperio AT2. Decrosslinking was carried out, followed by an overnight probe hybridization. Both left-hand and right-hand probes were ligated, and tissues were stained with eosin. The slide, equipped with the appropriate CytAssist Gene Expression slide, probe release, and tissue digestion master mix, was inserted into the CytAssist. A low-resolution image was captured by the CytAssist for fiducial frame alignment, and probes were transferred from the charged glass slide and tissue onto the Gene Expression slide. The probes underwent elongation and subsequent elution from the Gene Expression slide, followed by amplification and purification. The final index PCR cycles were determined via qPCR (Kapa; Agilent). The quality of the final library was evaluated using the LabChip GX Touch HT (Revvity), and libraries were quantified using Qubit. Pooled libraries were then subjected to paired-end sequencing as per the manufacturer’s protocol (Illumina NovaSeqXPlus). BclConvert software (Illumina) was employed to generate de-multiplexed Fastq files. The CytAssist and high-resolution images were aligned using the 10X Genomics Loupe Alignment software, and the SpaceRanger Pipeline (10X Genomics) was utilized to align reads and generate count matrices. This enabled us to evaluate the differential expression between spots at a resolution of 55 µm.

Spatial transcriptomics was performed on one high-strain (>15% interfragmentary strain) and one low-strain (<5% strain) week-3 callus. One representative FFPE callus from each group was used for spatial profiling; serial adjacent sections were processed for picrosirius red, Movat’s pentachrome, and IHC (SP7, SOX9, CD31, α-SMA) for orthogonal validation. Orthogonal support was strongest for collagen-rich fibrosis (picrosirius/Movat’s) and the osteoblast marker SP7. SOX9, CD31, and α-SMA showed lower transcript abundance, limiting cross-modality confirmation for those markers (Lower concordance for SOX9, CD31, and α-SMA is expected given 55 µm spot mixing, and known RNA-protein discordance for these markers). Because the spatial dataset includes one representative sample per condition, cohort-level comparisons primarily reflect within-sample variability and, despite orthogonal validation, may not be fully representative of larger datasets.

LoupeR was used to carefully manually select the desired region of interest within the samples, mainly focusing on callus and intramedullary canal 1-2 mm surrounding the fracture line. We processed count matrices from sequencing data using R (version 4.3.2). Quality assessment involved filtering out cells with low unique molecular identifiers (UMI) or gene expression. Analysis regions with < 500 UMIs or < 250 expressed genes were excluded. Additionally, genes not expressed in at least three Analysis regions were removed. We evaluated the impact of mitochondrial expression and cell cycle effects and adjusted for them as needed. The Seurat (version 4.4.0) (*59*) package was employed for post-processing, including normalization (SCTransform v2), integration, principal component analysis (PCA), uniform manifold approximation and projection (UMAP), and clustering (40 dimensions, clustering resolution of 1.4). Clusters were assigned to cell types using ScType (*60*) and a custom gene set for bone cells derived from the Panglao database (*61*). Manual validation based on highly expressed unique genes confirmed cell type assignments. Differential gene expression analysis of cell types between test groups (high and low-strain groups) utilized the Wilcoxon rank sum test with Bonferroni correction. Differentially expressed genes (DEGs) were filtered to only include genes with an average *log*_2_ fold change of > 0.25 or < -0.25 and adjusted p-value < 0.05. The filtered up and downregulated genes were then analyzed through gene ontology (GO) enrichment of biological processes using clusterProfiler (version 4.8.3), AnnotationDbi (version 1.62.2), and org.Mm.eg.db (version 3.17.0) packages. Finally, communication networks between cell types were explored using the CellChat (*62*) package (version 2.1.2).

### 7. Flow cytometry and multiplex assay

Blood samples were collected at specific time points and processed as described above. Briefly, plasma was separated through centrifugation, and red blood cells (RBC) were lysed from the remaining sample following the manufacturer’s guidelines using 1X RBC lysis buffer (eBioscience). Subsequently, an equal volume of 1X PBS (Gibco) was added to the suspension, followed by centrifugation at 400 g for 5 minutes. The resulting cell pellet was suspended in a blocking solution consisting of FACS buffer (1X PBS with 5% fetal bovine serum (Gibco) and 0.1% sodium azide) along with anti-mouse CD16/32 (eBioscience) for 10 minutes at 4°C to block Fc receptors.

The cells were then stained with fluorescently conjugated anti-mouse antibodies in FACS buffer for 45 minutes in the dark at 4°C. Detailed information regarding the fluorescent conjugates, clones, dilution factors, and suppliers for these antibodies can be found in **Supplementary Table S4**. The cell identification markers are listed in **Supplementary Table S5.** Following the staining procedure, the solution was removed through centrifugation at 400 g for 5 minutes, and the cells were subsequently resuspended in fresh FACS buffer. The samples were promptly subjected to analysis using a BD FACSymphony A3, and the data were processed using FlowJo software. Fluorescence-minus-one (FMO) controls for each antibody were employed to establish negative gates, ensuring a noise level of less than 1%.

To investigate the presence of inflammatory proteins, cytokines, and chemokines in the blood samples, a 32-analyte Milliplex MAP Mouse Cytokine/Chemokine Panel (Millipore, MCYTMAG-70K-PX32) was utilized. Initially, the plasma was isolated as previously described and then diluted in accordance with the manufacturer’s recommendations. Capture beads were added to each well of the plate, which was subsequently washed using a Hand-Held Magnetic Plate Washer (EPX-55555-000). Following the wash, samples, standards, and low and high controls were added to the wells in duplicate. The plate was then sealed with the provided plate sealer and lid, mixed at a speed of 600 rpm for 2 hours at room temperature, and was washed thrice using the provided wash buffer. A detection antibody mix was added, and the plate was resealed and mixed for 30 minutes at room temperature. The plate was then washed thrice, and Streptavidin-PE was added, resealed, and agitated for 30 minutes at ambient temperature and washed thrice. After adding the reading buffer and incubating for 5 minutes, the plate was processed on an xMAP INTELLIFLEX system with a 50 µL acquisition volume, a DD Gate ranging from 4000 to 13000, and a low PMT reporter gain setting.

### 8. Univariate and multivariate analysis

We conducted a comprehensive, multifaceted analysis to investigate the relationship between immune cell populations, biochemical factors in both blood and callus tissues, and healing outcomes through conventional linear regression. Additionally, we employed multivariate models, including linear and logistic regression. This included partial least squares regression (PLSR) analysis, which incorporated weighted averages of cell populations, cytokines, chemokines, and healing outcomes to identify the most influential immune factors. We utilized predictive analytics software JMP-PRO for all analyses. PLSR created principal components (latent variables, LV) combining immune cells, factors, and healing measures, and we assessed their predictive power by correlating them with the immune factors. We considered quadratic terms and variable interactions to enhance the model fits and employed the NIPALS algorithm with a 7 KFOLD validation method for generating linear plots. Notably, we focused on the first two linear combinations to identify the most influential healing factors.

### 9. Statistics

Data are plotted as either means with error bars representing the standard deviation or medians with error bars representing the interquartile range. The Pearson correlation coefficient (r) was used to evaluate the linear correlation between two variables, and the t-distribution was used to evaluate the associated p-value. To ensure the reliability of statistical comparisons, we subjected all groups to tests for normality and equal variance, employing the Shapiro-Wilk and Brown-Forsythe tests, respectively. In cases where both conditions were met, we used Student’s T-test for two independent groups. When normality was not met, we applied the Mann-Whitney Rank Sum Test, and if the equal variance test failed, a Welch’s t-test was used. For the analysis of systemic markers over time, a 2-way repeated measures ANOVA was performed. All spatial differential expression and enrichment results are spot-level with FDR correction as implemented in Seurat. Because the spatial dataset is from one representative sample from each group, we did not compute sample-level p-values for group comparisons from spatial data. Cohort-level inferences in this study are based on non-spatial assays with appropriate sample sizes and statistics. For each cytokine, we fit a two-way ANOVA with fixed effects Condition and Time and their interaction in R (v4.4.0) using the stats package; undetected values were treated as missing. All statistical tests were performed in JMP, R, or GraphPad Prism. Differences with *p* < 0.05 were considered statistically significant.

## F. Supplementary

**Fig. S1.**
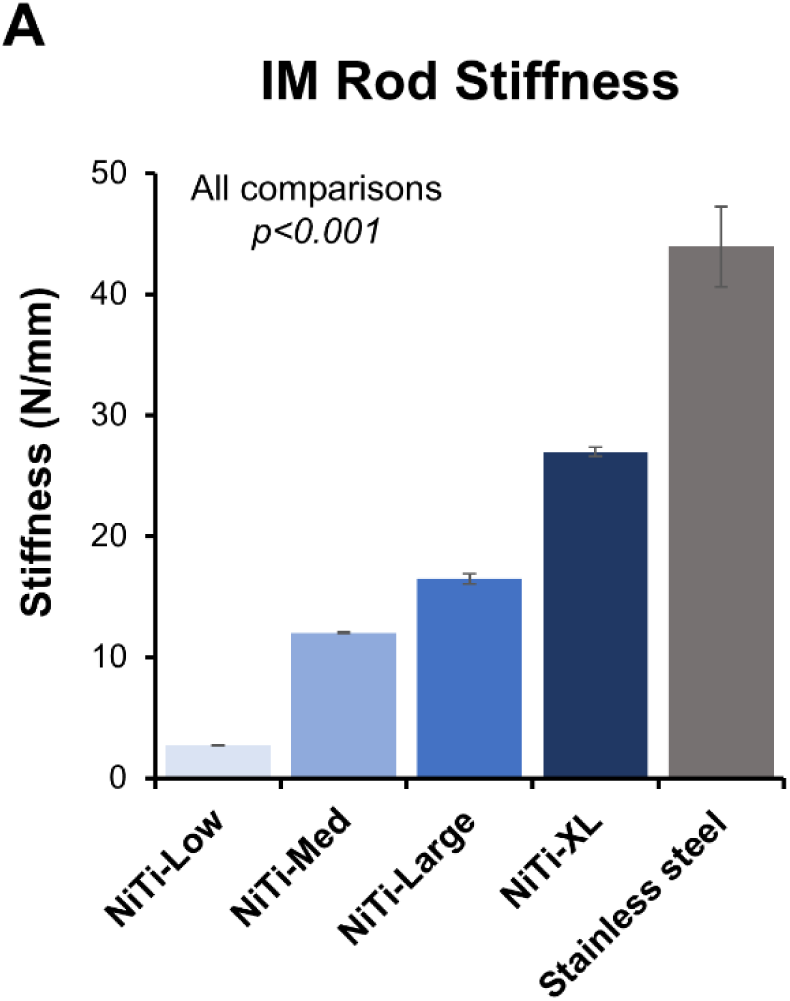
Tunable intramedullary (IM) rod stiffness across implant groups. Mechanical testing revealed progressive increases in axial stiffness across custom rod designs (Low, Medium, High, X-High) compared to a standard medical-grade 316L stainless steel nail. All group comparisons were statistically significant (*p* < 0.001). Error bars represent standard deviation.

**Fig. S2.**
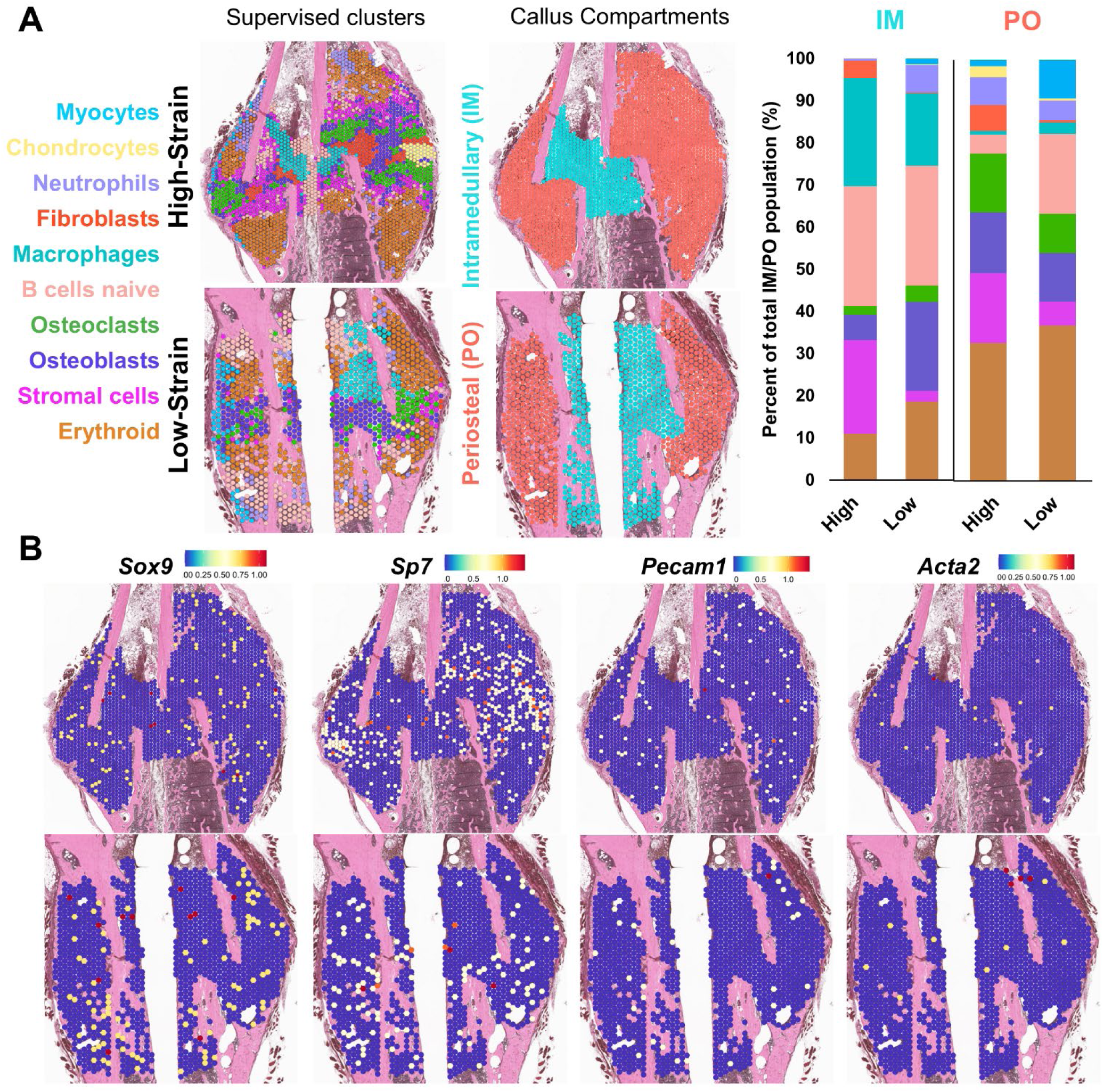
Spatial transcriptomic analysis of cell populations and IHC-associated markers. (**A**) Histological sections of high-strain (low-stiffness nail, >15% interfragmentary strain, top) and low-strain (high-stiffness nail, <5% interfragmentary strain, bottom) calluses overlaid with supervised spatial transcriptomic clusters (left) and manually defined callus compartments (middle), with corresponding cell population percentages in periosteal (PO) and intramedullary (IM) regions (right). (**B**) Spatial expression plots of Sox9 (SOX9), Sp7 (SP7/Osterix), Pecam1 (CD31), and Acta2 (αSMA), the markers evaluated by immunohistochemistry. One representative FFPE callus from each group was used for spatial profiling; serial adjacent sections were processed for picrosirius red, Movat’s pentachrome, and IHC for orthogonal validation.

**Fig. S3.**
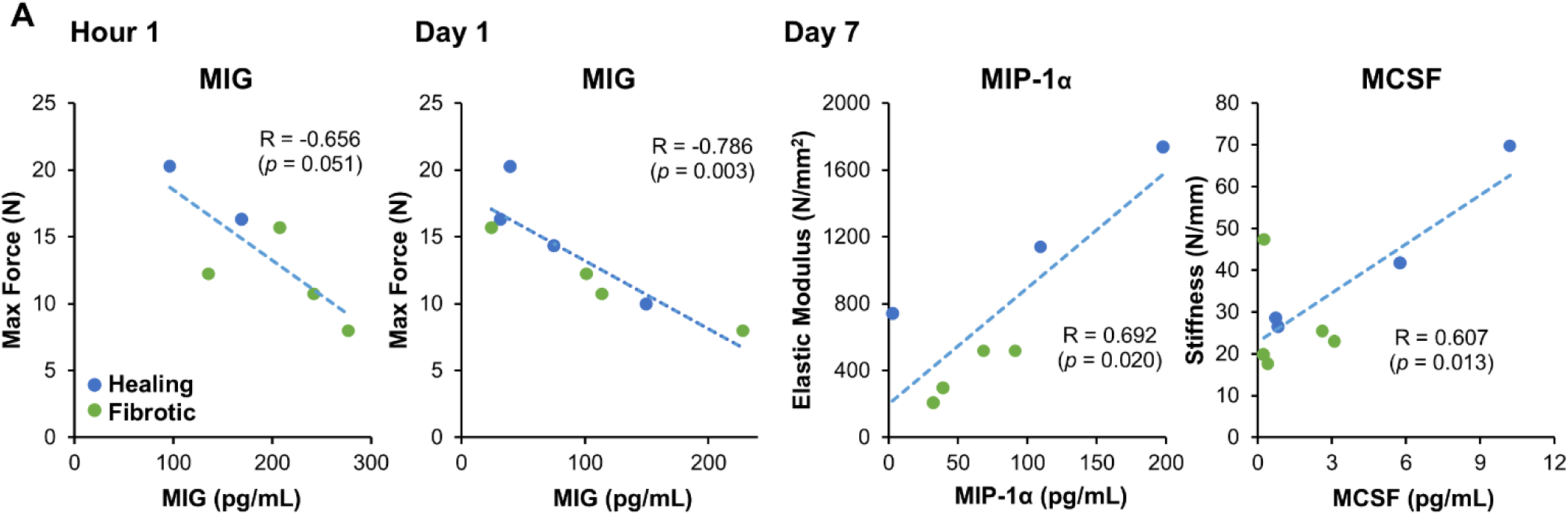
Cytokine and Chemokines Predict Biomechanical Healing Outcomes. (**A**) Hour 1 post-fracture: Negative correlation between MIG and maximum force (R = -0.656, p = 0.051). Day 1 post-fracture: Negative correlation between MIG and maximum force (R = -0.786, p = 0.003). Day 7 post-fracture: Positive correlation between MIP-1α and elastic modulus (R = 0.692, p = 0.020), and a positive correlation between MCSF and stiffness (R = 0.607, p = 0.013). Data are linear regression fits, n = 7 per group displaying only detected values. Green dots represent individual high-strain (low-stiffness nail, >15% interfragmentary strain) samples, blue dots represent individual low-strain (high-stiffness nail, <5% interfragmentary strain) samples.

**Table S1.**
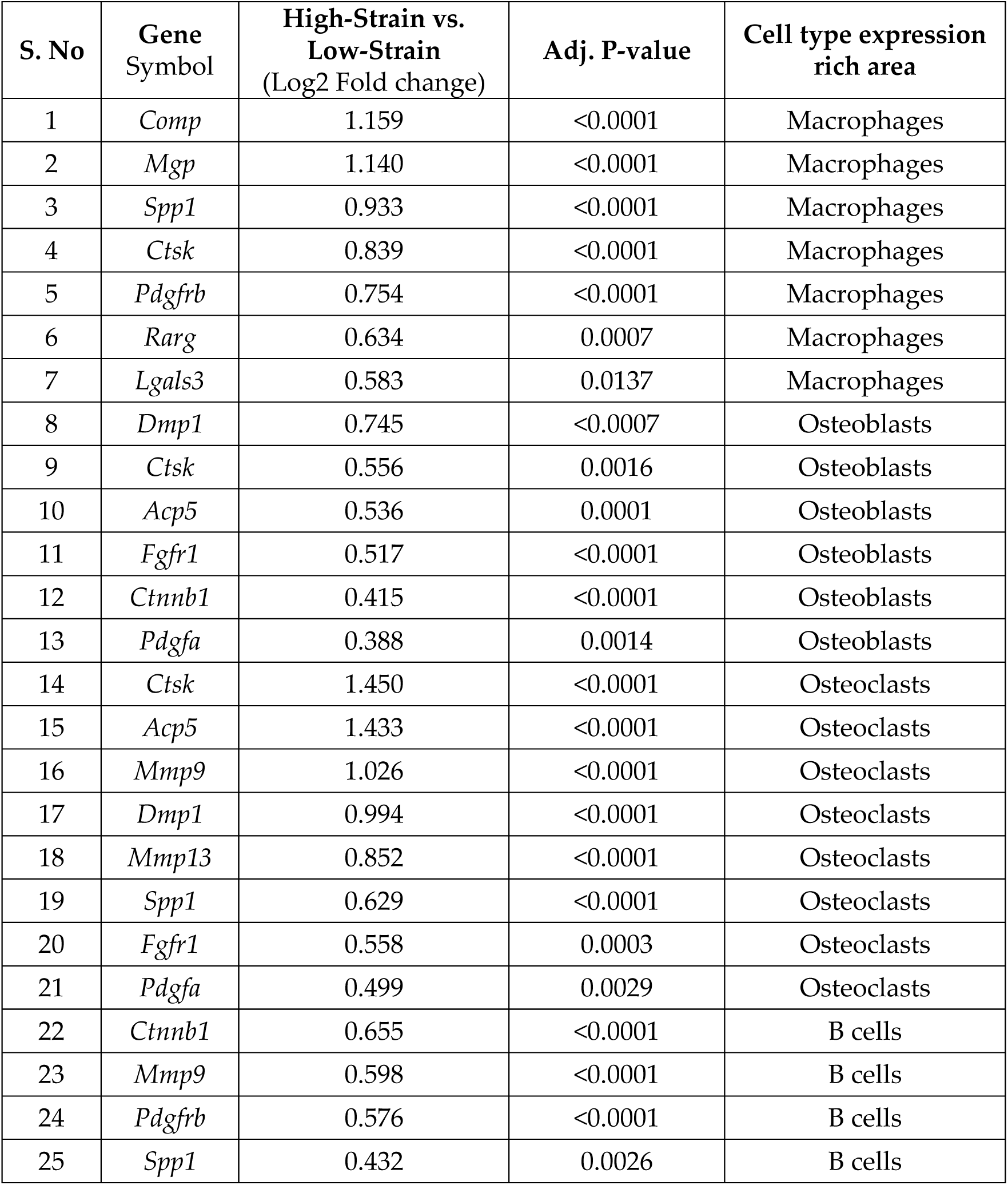
Spatial Transcriptomics Fibrotic Expression Analysis. Significant differentially expressed fibrotic genes between the high-strain (low-stiffness nail) and low-strain (high-stiffness nail) groups. The values presented are log2 Fold changes.

**Table S2.**
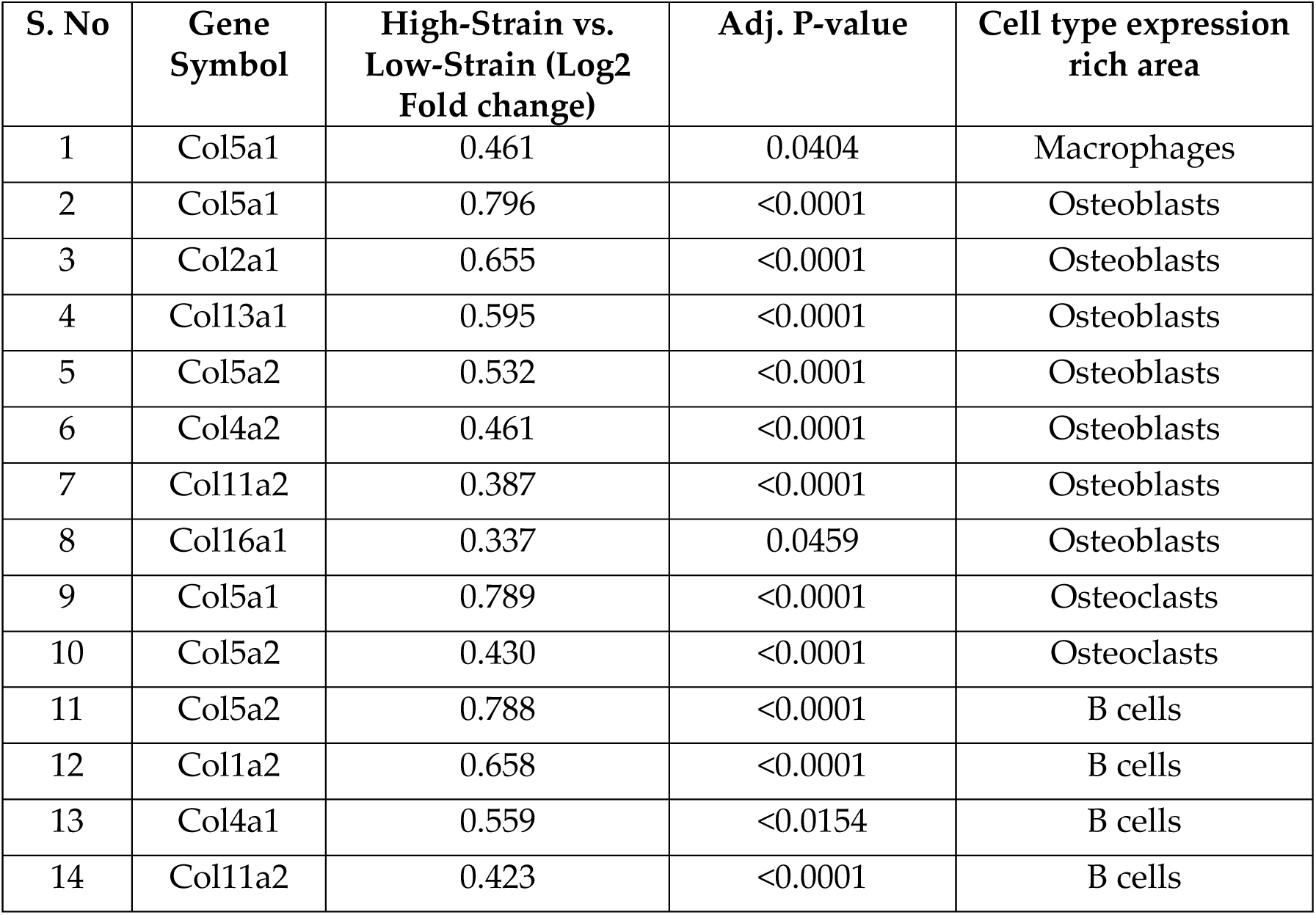
Spatial Transcriptomics Collagen Expression Analysis. Significant differentially expressed collagen genes between the high-strain (low-stiffness nail) and low-strain (high-stiffness nail) groups. The values presented are log2 Fold changes.

**Table S3.**
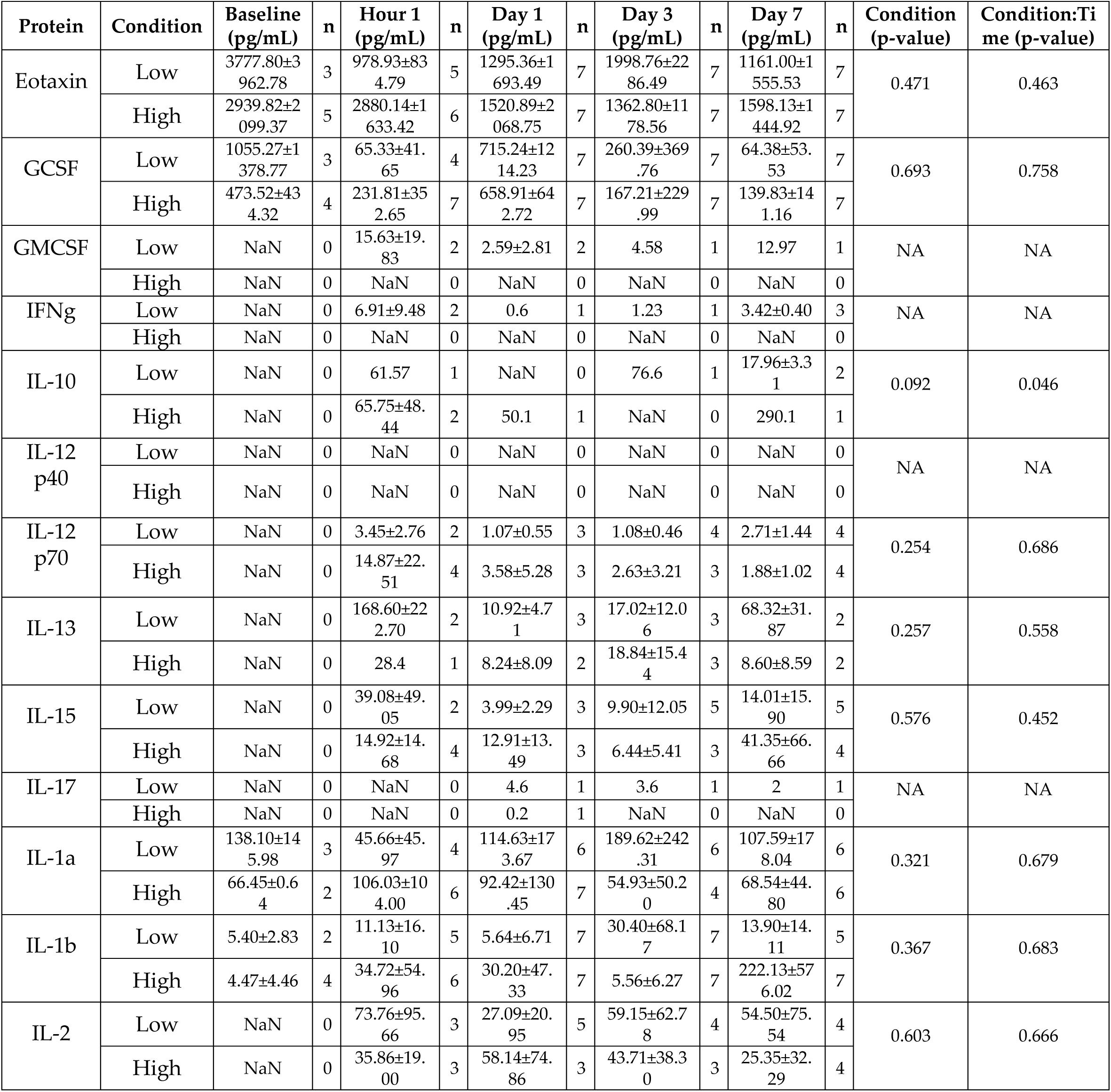
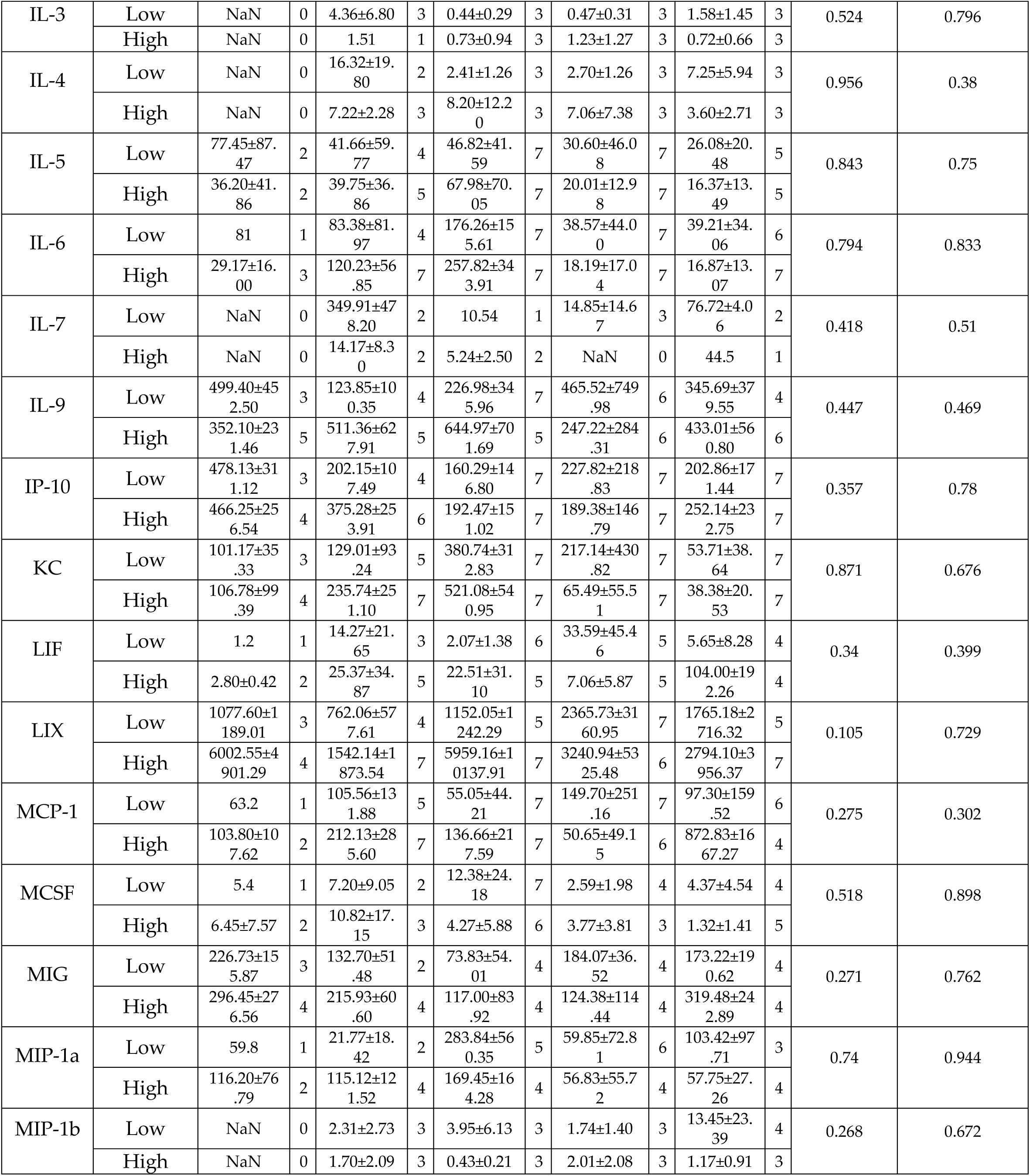
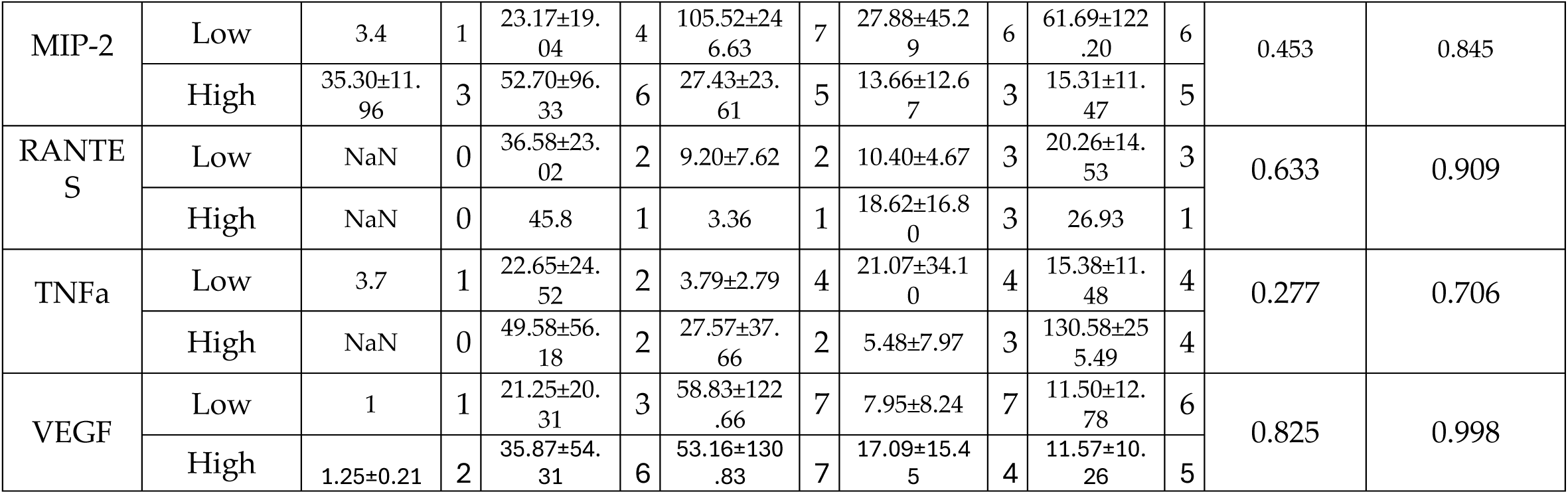
Systemic Plasma Cytokines after Fracture. Mean ± SD (pg/mL) at Baseline, 1 h, Day 1, Day 3, and Day 7 in Low-strain (high-stiffness nail) vs. High-strain (low-stiffness nail) groups. n gives samples per time point. P-values are from a two-way ANOVA: P-values for Condition and Condition:Time are reported for each protein. NaN indicates no measurable signal (protein not detected); NA indicates insufficient data to fit the ANOVA.

| Protein | Condition | Baseline (pg/mL) | n | Hour 1 (pg/mL) | n | Day 1 (pg/mL) | n | Day 3 (pg/mL) | n | Day 7 (pg/mL) | n | Condition (p-value) | Condition:Time (p-value) |
| --- | --- | --- | --- | --- | --- | --- | --- | --- | --- | --- | --- | --- | --- |
| Eotaxin | Low | 3777.80±3962.78 | 3 | 978.93±834.79 | 5 | 1295.36±1693.49 | 7 | 1998.76±2286.49 | 7 | 1161.00±1555.53 | 7 | 0.471 | 0.463 |
| | High | 2939.82±2099.37 | 5 | 2880.14±1633.42 | 6 | 1520.89±2068.75 | 7 | 1362.80±1178.56 | 7 | 1598.13±1444.92 | 7 | | |
| GCSF | Low | 1055.27±1378.77 | 3 | 65.33±41.65 | 4 | 715.24±1214.23 | 7 | 260.39±369.76 | 7 | 64.38±53.53 | 7 | 0.693 | 0.758 |
| | High | 473.52±434.32 | 4 | 231.81±352.65 | 7 | 658.91±642.72 | 7 | 167.21±229.99 | 7 | 139.83±141.16 | 7 | | |
| GMCSF | Low | NaN | 0 | 15.63±19.83 | 2 | 2.59±2.81 | 2 | 4.58 | 1 | 12.97 | 1 | NA | NA |
|  | High | NaN | 0 | NaN | 0 | NaN | 0 | NaN | 0 | NaN | 0 |  |  |
| IFNg | Low | NaN | 0 | 6.91±9.48 | 2 | 0.6 | 1 | 1.23 | 1 | 3.42±0.40 | 3 | NA | NA |
|  | High | NaN | 0 | NaN | 0 | NaN | 0 | NaN | 0 | NaN | 0 |  |  |
| IL-10 | Low | NaN | 0 | 61.57 | 1 | NaN | 0 | 76.6 | 1 | 17.96±3.31 | 2 | 0.092 | 0.046 |
| | High | NaN | 0 | 65.75±48.44 | 2 | 50.1 | 1 | NaN | 0 | 290.1 | 1 | | |
| IL-12 p40 | Low | NaN | 0 | NaN | 0 | NaN | 0 | NaN | 0 | NaN | 0 | NA | NA |
|  | High | NaN | 0 | NaN | 0 | NaN | 0 | NaN | 0 | NaN | 0 |  |  |
| IL-12 p70 | Low | NaN | 0 | 3.45±2.76 | 2 | 1.07±0.55 | 3 | 1.08±0.46 | 4 | 2.71±1.44 | 4 | 0.254 | 0.686 |
| | High | NaN | 0 | 14.87±22.51 | 4 | 3.58±5.28 | 3 | 2.63±3.21 | 3 | 1.88±1.02 | 4 | | |
| IL-13 | Low | NaN | 0 | 168.60±222.70 | 2 | 10.92±4.71 | 3 | 17.02±12.06 | 3 | 68.32±31.87 | 2 | 0.257 | 0.558 |
| | High | NaN | 0 | 28.4 | 1 | 8.24±8.09 | 2 | 18.84±15.44 | 3 | 8.60±8.59 | 2 | | |
| IL-15 | Low | NaN | 0 | 39.08±49.05 | 2 | 3.99±2.29 | 3 | 9.90±12.05 | 5 | 14.01±15.90 | 5 | 0.576 | 0.452 |
| | High | NaN | 0 | 14.92±14.68 | 4 | 12.91±13.49 | 3 | 6.44±5.41 | 3 | 41.35±66.66 | 4 | | |
| IL-17 | Low | NaN | 0 | NaN | 0 | 4.6 | 1 | 3.6 | 1 | 2 | 1 | NA | NA |
|  | High | NaN | 0 | NaN | 0 | 0.2 | 1 | NaN | 0 | NaN | 0 |  |  |
| IL-1a | Low | 138.10±145.98 | 3 | 45.66±45.97 | 4 | 114.63±173.67 | 6 | 189.62±242.31 | 6 | 107.59±178.04 | 6 | 0.321 | 0.679 |
| | High | 66.45±0.64 | 2 | 106.03±104.00 | 6 | 92.42±130.45 | 7 | 54.93±50.20 | 4 | 68.54±44.80 | 6 | | |
| IL-1b | Low | 5.40±2.83 | 2 | 11.13±16.10 | 5 | 5.64±6.71 | 7 | 30.40±68.17 | 7 | 13.90±14.11 | 5 | 0.367 | 0.683 |
| | High | 4.47±4.46 | 4 | 34.72±54.96 | 6 | 30.20±47.33 | 7 | 5.56±6.27 | 7 | 222.13±576.02 | 7 | | |
| IL-2 | Low | NaN | 0 | 73.76±95.66 | 3 | 27.09±20.95 | 5 | 59.15±62.78 | 4 | 54.50±75.54 | 4 | 0.603 | 0.666 |
| | High | NaN | 0 | 35.86±19.00 | 3 | 58.14±74.86 | 3 | 43.71±38.30 | 3 | 25.35±32.29 | 4 | | |
| IL-3 | Low | NaN | 0 | 4.36±6.80 | 3 | 0.44±0.29 | 3 | 0.47±0.31 | 3 | 1.58±1.45 | 3 | 0.524 | 0.796 |
|  | High | NaN | 0 | 1.51 | 1 | 0.73±0.94 | 3 | 1.23±1.27 | 3 | 0.72±0.66 | 3 |  |  |
| IL-4 | Low | NaN | 0 | 16.32±19.80 | 2 | 2.41±1.26 | 3 | 2.70±1.26 | 3 | 7.25±5.94 | 3 | 0.956 | 0.38 |
|  | High | NaN | 0 | 7.22±2.28 | 3 | 8.20±12.20 | 3 | 7.06±7.38 | 3 | 3.60±2.71 | 3 |  |  |
| IL-5 | Low | 77.45±87.47 | 2 | 41.66±59.77 | 4 | 46.82±41.59 | 7 | 30.60±46.08 | 7 | 26.08±20.48 | 5 | 0.843 | 0.75 |
|  | High | 36.20±41.86 | 2 | 39.75±36.86 | 5 | 67.98±70.05 | 7 | 20.01±12.98 | 7 | 16.37±13.49 | 5 |  |  |
| IL-6 | Low | 81 | 1 | 83.38±81.97 | 4 | 176.26±155.61 | 7 | 38.57±44.00 | 7 | 39.21±34.06 | 6 | 0.794 | 0.833 |
|  | High | 29.17±16.00 | 3 | 120.23±56.85 | 7 | 257.82±343.91 | 7 | 18.19±17.04 | 7 | 16.87±13.07 | 7 |  |  |
| IL-7 | Low | NaN | 0 | 349.91±478.20 | 2 | 10.54 | 1 | 14.85±14.67 | 3 | 76.72±4.06 | 2 | 0.418 | 0.51 |
|  | High | NaN | 0 | 14.17±8.30 | 2 | 5.24±2.50 | 2 | NaN | 0 | 44.5 | 1 |  |  |
| IL-9 | Low | 499.40±452.50 | 3 | 123.85±100.35 | 4 | 226.98±345.96 | 7 | 465.52±749.98 | 6 | 345.69±379.55 | 4 | 0.447 | 0.469 |
|  | High | 352.10±231.46 | 5 | 511.36±627.91 | 5 | 644.97±701.69 | 5 | 247.22±284.31 | 6 | 433.01±560.80 | 6 |  |  |
| IP-10 | Low | 478.13±311.12 | 3 | 202.15±107.49 | 4 | 160.29±146.80 | 7 | 227.82±218.83 | 7 | 202.86±171.44 | 7 | 0.357 | 0.78 |
|  | High | 466.25±256.54 | 4 | 375.28±253.91 | 6 | 192.47±151.02 | 7 | 189.38±146.79 | 7 | 252.14±232.75 | 7 |  |  |
| KC | Low | 101.17±35.33 | 3 | 129.01±93.24 | 5 | 380.74±312.83 | 7 | 217.14±430.82 | 7 | 53.71±38.64 | 7 | 0.871 | 0.676 |
|  | High | 106.78±99.39 | 4 | 235.74±251.10 | 7 | 521.08±540.95 | 7 | 65.49±55.51 | 7 | 38.38±20.53 | 7 |  |  |
| LIF | Low | 1.2 | 1 | 14.27±21.65 | 3 | 2.07±1.38 | 6 | 33.59±45.46 | 5 | 5.65±8.28 | 4 | 0.34 | 0.399 |
|  | High | 2.80±0.42 | 2 | 25.37±34.87 | 5 | 22.51±31.10 | 5 | 7.06±5.87 | 5 | 104.00±192.26 | 4 |  |  |
| LIX | Low | 1077.60±1189.01 | 3 | 762.06±577.61 | 4 | 1152.05±1242.29 | 5 | 2365.73±3160.95 | 7 | 1765.18±2716.32 | 5 | 0.105 | 0.729 |
|  | High | 6002.55±4901.29 | 4 | 1542.14±1873.54 | 7 | 5959.16±10137.91 | 7 | 3240.94±5325.48 | 6 | 2794.10±3956.37 | 7 |  |  |
| MCP-1 | Low | 63.2 | 1 | 105.56±131.88 | 5 | 55.05±44.21 | 7 | 149.70±251.16 | 7 | 97.30±159.52 | 6 | 0.275 | 0.302 |
|  | High | 103.80±107.62 | 2 | 212.13±285.60 | 7 | 136.66±217.59 | 7 | 50.65±49.15 | 6 | 872.83±1667.27 | 4 |  |  |
| MCSF | Low | 5.4 | 1 | 7.20±9.05 | 2 | 12.38±24.18 | 7 | 2.59±1.98 | 4 | 4.37±4.54 | 4 | 0.518 | 0.898 |
|  | High | 6.45±7.57 | 2 | 10.82±17.15 | 3 | 4.27±5.88 | 6 | 3.77±3.81 | 3 | 1.32±1.41 | 5 |  |  |
| MIG | Low | 226.73±155.87 | 3 | 132.70±51.48 | 2 | 73.83±54.01 | 4 | 184.07±36.52 | 4 | 173.22±190.62 | 4 | 0.271 | 0.762 |
|  | High | 296.45±276.56 | 4 | 215.93±60.60 | 4 | 117.00±83.92 | 4 | 124.38±114.44 | 4 | 319.48±242.89 | 4 |  |  |
| MIP-1a | Low | 59.8 | 1 | 21.77±18.42 | 2 | 283.84±560.35 | 5 | 59.85±72.81 | 6 | 103.42±97.71 | 3 | 0.74 | 0.944 |
|  | High | 116.20±76.79 | 2 | 115.12±121.52 | 4 | 169.45±164.28 | 4 | 56.83±55.72 | 4 | 57.75±27.26 | 4 |  |  |
| MIP-1b | Low | NaN | 0 | 2.31±2.73 | 3 | 3.95±6.13 | 3 | 1.74±1.40 | 3 | 13.45±23.39 | 4 | 0.268 | 0.672 |
|  | High | NaN | 0 | 1.70±2.09 | 3 | 0.43±0.21 | 3 | 2.01±2.08 | 3 | 1.17±0.91 | 3 |  |  |
| MIP-2 | Low | 3.4 | 1 | 23.17±19.04 | 4 | 105.52±246.63 | 7 | 27.88±45.29 | 6 | 61.69±122.20 | 6 | 0.453 | 0.845 |
|  | High | 35.30±11.96 | 3 | 52.70±96.33 | 6 | 27.43±23.61 | 5 | 13.66±12.67 | 3 | 15.31±11.47 | 5 |  |  |
| RANTES | Low | NaN | 0 | 36.58±23.02 | 2 | 9.20±7.62 | 2 | 10.40±4.67 | 3 | 20.26±14.53 | 3 | 0.633 | 0.909 |
|  | High | NaN | 0 | 45.8 | 1 | 3.36 | 1 | 18.62±16.80 | 3 | 26.93 | 1 |  |  |
| TNFa | Low | 3.7 | 1 | 22.65±24.52 | 2 | 3.79±2.79 | 4 | 21.07±34.10 | 4 | 15.38±11.48 | 4 | 0.277 | 0.706 |
|  | High | NaN | 0 | 49.58±56.18 | 2 | 27.57±37.66 | 2 | 5.48±7.97 | 3 | 130.58±255.49 | 4 |  |  |
| VEGF | Low | 1 | 1 | 21.25±20.31 | 3 | 58.83±122.66 | 7 | 7.95±8.24 | 7 | 11.50±12.78 | 6 | 0.825 | 0.998 |
|  | High | 1.25±0.21 | 2 | 35.87±54.31 | 6 | 53.16±130.83 | 7 | 17.09±15.45 | 4 | 11.57±10.26 | 5 |  |  |

**Table S4.**
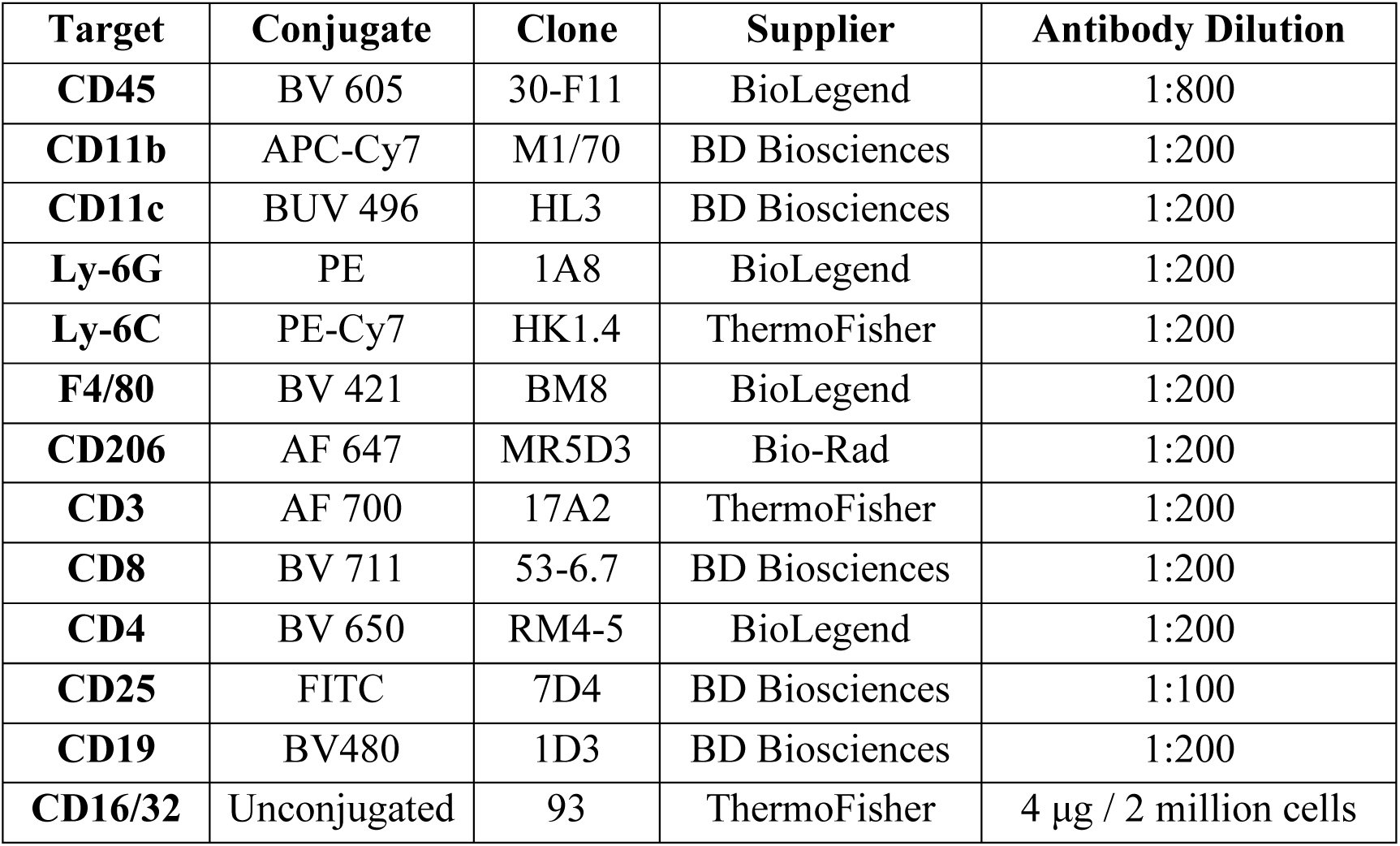
Flow Cytometry Antibody Information. List of surface markers, fluorophore conjugates, clones, suppliers, and working dilutions used for staining murine immune cell populations. All antibodies were used at the specified dilutions, and Fc receptors were blocked with anti-CD16/32 (clone 93) at 4 µg per 2 million cells to prevent nonspecific binding.

**Table S5.**
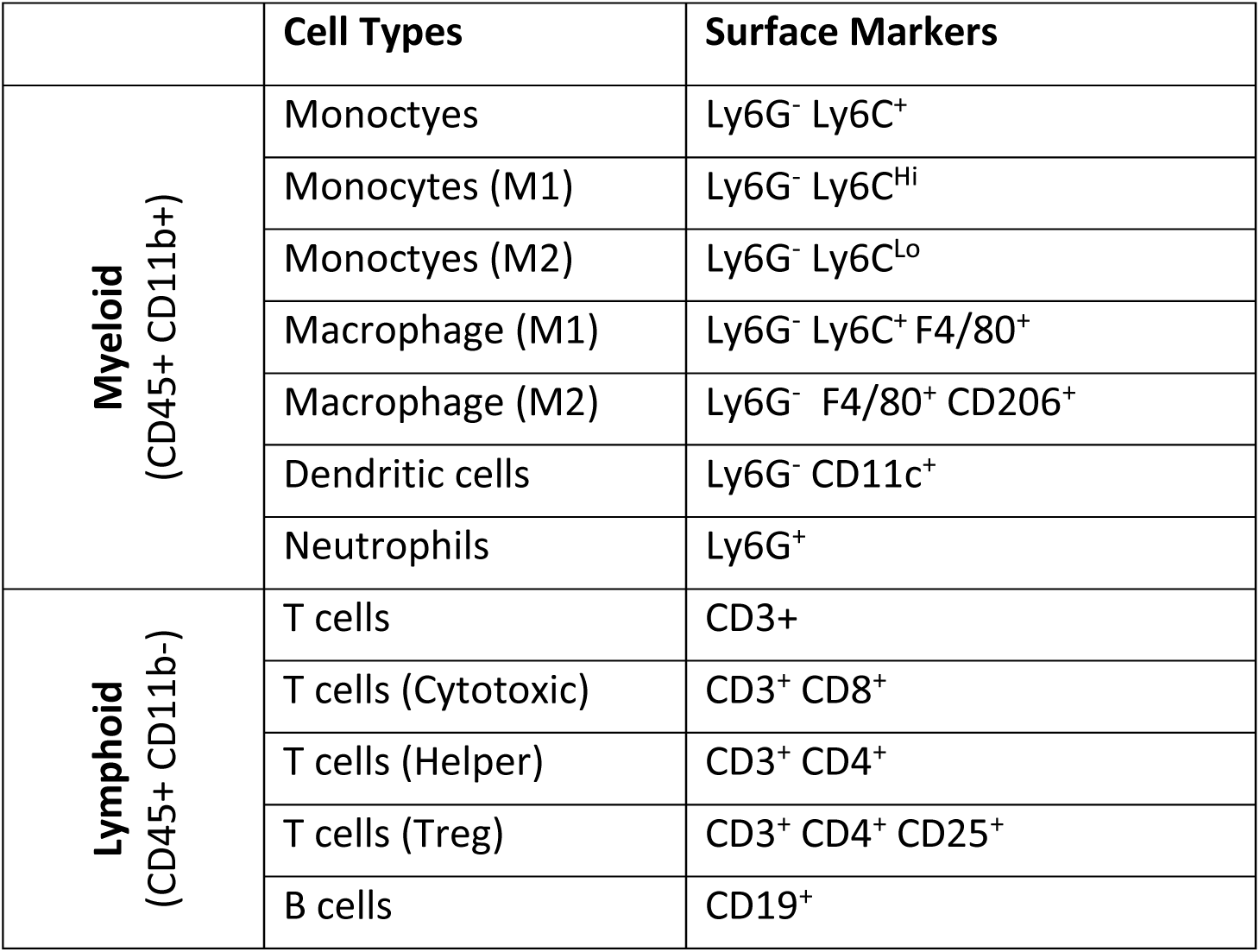
Flow Cytometry Cell Type Surface Markers. Immune cell populations were identified from CD45⁺ cells and subdivided into myeloid (CD11b⁺) and lymphoid (CD11b⁻) lineages. Subsets were classified using the cell surface markers below.

## Acknowledgements

We thank the Advanced Genomics Core at the University of Michigan for library prep and next-generation sequencing. We gratefully acknowledge the late Dr. Arnold Stromberg (Department of Statistics, University of Kentucky) for his invaluable contributions to statistical modeling and analysis.

## Competing interests

All authors declare that they have no competing interests.

## Funding

Research reported in this publication was supported in part by the National Institute of Arthritis and Musculoskeletal and Skin Diseases (NIAMS) Award Numbers R21AR078447 and R01AR066028, National Institute of General Medical Sciences (NIGMS) of the National Institutes of Health under Award Number P20GM130456, Orthopedic Trauma Association (OTA, Grant Number: 6889), and Biospecimen Procurement & Translational Pathology Shared Resource Facility of the University of Kentucky Markey Cancer Center Award Number P30CA177558. The content is solely the responsibility of the authors and does not necessarily represent the official views of the National Institutes of Health or other grant funding agencies.

## Author contributions

Conceptualization: R.T.A., M.D.P., K.D.H., and J.A. Methodology: R.T.A., M.D.P., K.D.H., and J.A. Software: R.T.A. and M.D.P. Validation: R.T.A. and M.D.P. Formal analysis: R.T.A. and M.D.P. Investigation: R.T.A. and M.D.P. Resources: R.T.A. and M.D.P. Data curation: R.T.A. and M.D.P. Writing - original draft: R.T.A. and M.D.P. Writing - review and editing: R.T.A., M.D.P., K.D.H., and J.A. Visualization: R.T.A. and M.D.P. Supervision: R.T.A., K.D.H., and J.A. Project administration: R.T.A. Funding Acquisition: R.T.A. and K.D.H.

## Data availability statement

All data needed to evaluate and reproduce the conclusions in the paper are present in the paper and/or the Supplementary Materials.

## Notes

### Competing Interest Statement

The authors have declared no competing interest.

### Summary of Updates

Revised the manuscript to update the text, incorporate new data and figures, and revise the Discussion section

