## Supplementary material for "Maladaptive immune-fibrotic axis drives impaired long bone regeneration under mechanical instability": Supplemetary data

Matthew D. Patrick *et al.*

**This PDF file includes:**

Fig. S1 to S3
Tables S1 to S5

**Fig. S1. Tunable intramedullary (IM) rod stiffness across implant groups.**
Mechanical testing revealed progressive increases in axial stiffness across custom rod designs (Low, Medium, High, X-High) compared to a standard medical-grade 316L stainless steel nail. All group comparisons were statistically significant ($p < 0.001$). Error bars represent standard deviation.

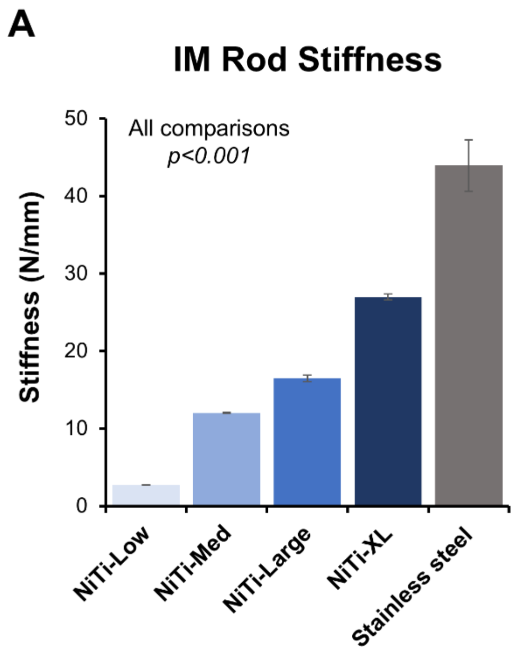

**Fig. S2. Spatial transcriptomic analysis of cell populations and IHC-associated markers.** (**A**) Histological sections of high-strain (low-stiffness nail, >15% interfragmentary strain, top) and low-strain (high-stiffness nail, <5% interfragmentary strain, bottom) calluses overlaid with supervised spatial transcriptomic clusters (left) and manually defined callus compartments (middle), with corresponding cell population percentages in periosteal (PO) and intramedullary (IM) regions (right). (**B**) Spatial expression plots of Sox9 (SOX9), Sp7 (SP7/Osterix), Pecam1 (CD31), and Acta2 (αSMA), the markers evaluated by immunohistochemistry. One representative FFPE callus from each group was used for spatial profiling; serial adjacent sections were processed for picrosirius red, Movat's pentachrome, and IHC for orthogonal validation.

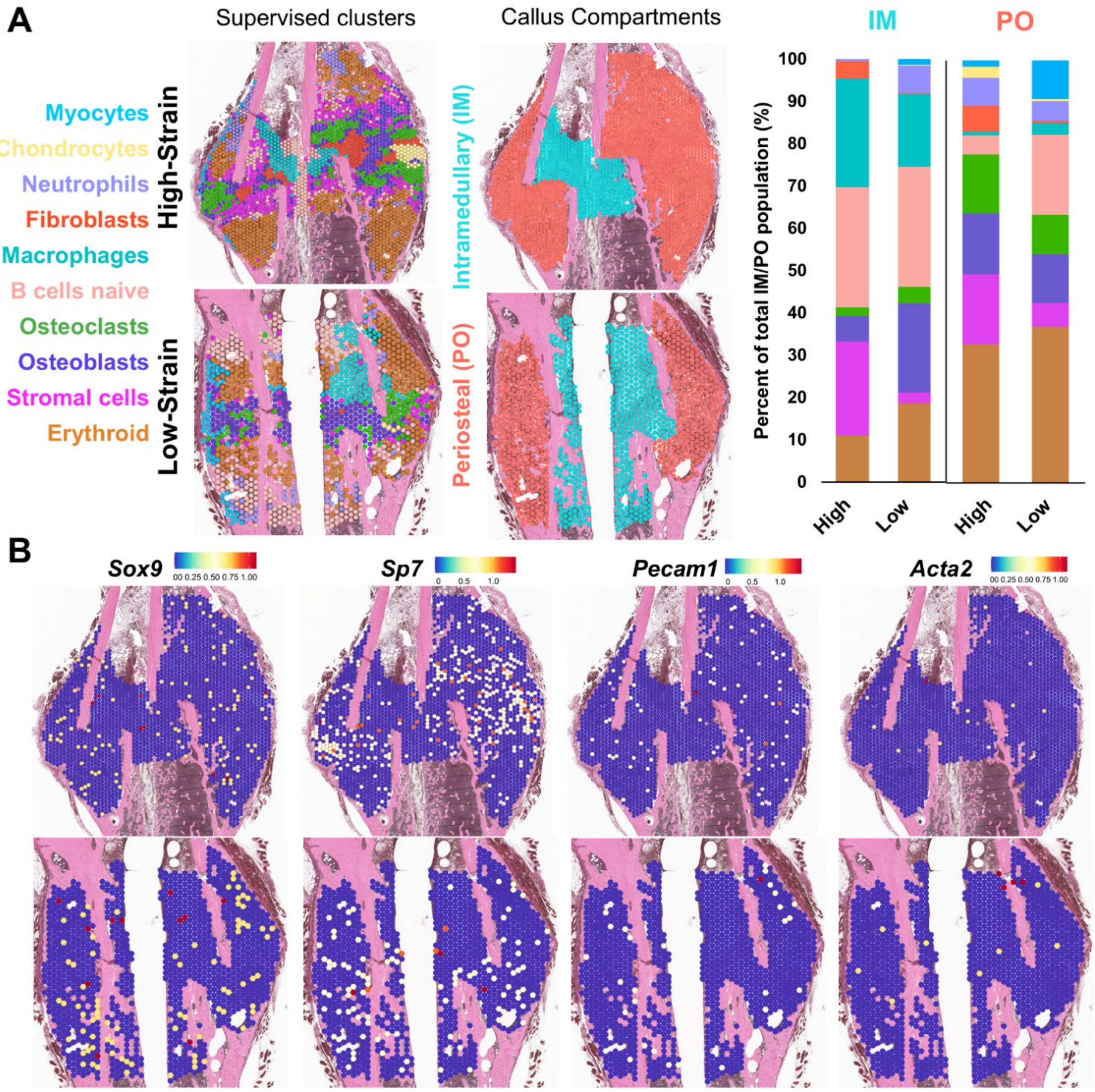

**Fig. S3. Cytokine and Chemokines Predict Biomechanical Healing Outcomes.**

(**A**) Hour 1 post-fracture: Negative correlation between MIG and maximum force (R = -0.656, p = 0.051). Day 1 post-fracture: Negative correlation between MIG and maximum force (R = -0.786, p = 0.003). Day 7 post-fracture: Positive correlation between MIP-1α and elastic modulus (R = 0.692, p = 0.020), and a positive correlation between MCSF and stiffness (R = 0.607, p = 0.013). Data are linear regression fits, n = 7 per group displaying only detected values. Green dots represent individual high-strain (low-stiffness nail, >15% interfragmentary strain) samples, blue dots represent individual low-strain (high-stiffness nail, <5% interfragmentary strain) samples.

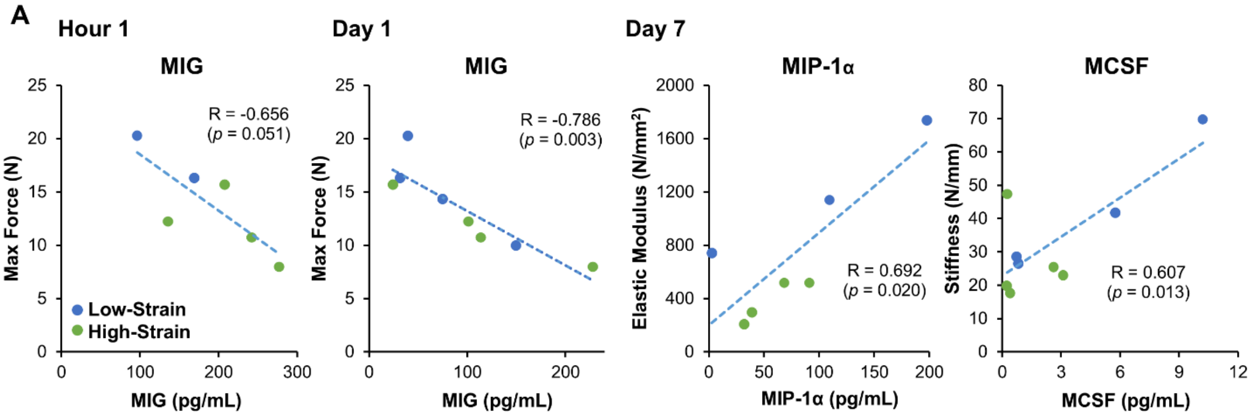

**Table S1. Spatial Transcriptomics Fibrotic Expression Analysis.**

*Significant differentially expressed fibrotic genes between the high-strain (low-stiffness nail) and low-strain (high-stiffness nail) groups.* The values presented are log2 Fold changes.

| S. No | Gene Symbol | High-Strain vs. Low-Strain (Log2 Fold change) | Adj. P-value | Cell type expression rich area |
|---|---|---|---|---|
| 1 | *Comp* | 1.159 | <0.0001 | Macrophages |
| 2 | *Mgp* | 1.140 | <0.0001 | Macrophages |
| 3 | *Spp1* | 0.933 | <0.0001 | Macrophages |
| 4 | *Ctsk* | 0.839 | <0.0001 | Macrophages |
| 5 | *Pdgfrb* | 0.754 | <0.0001 | Macrophages |
| 6 | *Rarg* | 0.634 | 0.0007 | Macrophages |
| 7 | *Lgals3* | 0.583 | 0.0137 | Macrophages |
| 8 | *Dmp1* | 0.745 | <0.0007 | Osteoblasts |
| 9 | *Ctsk* | 0.556 | 0.0016 | Osteoblasts |
| 10 | *Acp5* | 0.536 | 0.0001 | Osteoblasts |
| 11 | *Fgfr1* | 0.517 | <0.0001 | Osteoblasts |
| 12 | *Ctnnb1* | 0.415 | <0.0001 | Osteoblasts |
| 13 | *Pdgfa* | 0.388 | 0.0014 | Osteoblasts |
| 14 | *Ctsk* | 1.450 | <0.0001 | Osteoclasts |
| 15 | *Acp5* | 1.433 | <0.0001 | Osteoclasts |
| 16 | *Mmp9* | 1.026 | <0.0001 | Osteoclasts |
| 17 | *Dmp1* | 0.994 | <0.0001 | Osteoclasts |
| 18 | *Mmp13* | 0.852 | <0.0001 | Osteoclasts |
| 19 | *Spp1* | 0.629 | <0.0001 | Osteoclasts |
| 20 | *Fgfr1* | 0.558 | 0.0003 | Osteoclasts |
| 21 | *Pdgfa* | 0.499 | 0.0029 | Osteoclasts |
| 22 | *Ctnnb1* | 0.655 | <0.0001 | B cells |
| 23 | *Mmp9* | 0.598 | <0.0001 | B cells |
| 24 | *Pdgfrb* | 0.576 | <0.0001 | B cells |
| 25 | *Spp1* | 0.432 | 0.0026 | B cells |

**Table S2. Spatial Transcriptomics Collagen Expression Analysis.**

Significant differentially expressed collagen genes between the high-strain (low-stiffness nail) and low-strain (high-stiffness nail) groups. The values presented are log2 Fold changes.

| S. No | Gene Symbol | High-Strain vs. Low-Strain (Log2 Fold change) | Adj. P-value | Cell type expression rich area |
|---|---|---|---|---|
| 1 | Col5a1 | 0.461 | 0.0404 | Macrophages |
| 2 | Col5a1 | 0.796 | <0.0001 | Osteoblasts |
| 3 | Col2a1 | 0.655 | <0.0001 | Osteoblasts |
| 4 | Col13a1 | 0.595 | <0.0001 | Osteoblasts |
| 5 | Col5a2 | 0.532 | <0.0001 | Osteoblasts |
| 6 | Col4a2 | 0.461 | <0.0001 | Osteoblasts |
| 7 | Col11a2 | 0.387 | <0.0001 | Osteoblasts |
| 8 | Col16a1 | 0.337 | 0.0459 | Osteoblasts |
| 9 | Col5a1 | 0.789 | <0.0001 | Osteoclasts |
| 10 | Col5a2 | 0.430 | <0.0001 | Osteoclasts |
| 11 | Col5a2 | 0.788 | <0.0001 | B cells |
| 12 | Col1a2 | 0.658 | <0.0001 | B cells |
| 13 | Col4a1 | 0.559 | <0.0154 | B cells |
| 14 | Col11a2 | 0.423 | <0.0001 | B cells |

**Table S3. Systemic Plasma Cytokines after Fracture.**

Mean ± SD (pg/mL) at Baseline, 1 h, Day 1, Day 3, and Day 7 in Low-strain (high-stiffness nail) vs. High-strain (low-stiffness nail) groups. n gives samples per time point. P-values are from a two-way ANOVA: P-values for Condition and Condition:Time are reported for each protein. NaN indicates no measurable signal (protein not detected); NA indicates insufficient data to fit the ANOVA.

| Protein | Condition | Baseline (pg/mL) | n | Hour 1 (pg/mL) | n | Day 1 (pg/mL) | n | Day 3 (pg/mL) | n | Day 7 (pg/mL) | n | Condition (p-value) | Condition:Time (p-value) |
|---|---|---|---|---|---|---|---|---|---|---|---|---|---|
| Eotaxin | Low | 3777.80±3962.78 | 3 | 978.93±834.79 | 5 | 1295.36±1693.49 | 7 | 1998.76±2286.49 | 7 | 1161.00±1555.53 | 7 | 0.471 | 0.463 |
| | High | 2939.82±2099.37 | 5 | 2880.14±1633.42 | 6 | 1520.89±2068.75 | 7 | 1362.80±1178.56 | 7 | 1598.13±1444.92 | 7 | | |
| GCSF | Low | 1055.27±1378.77 | 3 | 65.33±41.65 | 4 | 715.24±1214.23 | 7 | 260.39±369.76 | 7 | 64.38±53.53 | 7 | 0.693 | 0.758 |
| | High | 473.52±434.32 | 4 | 231.81±352.65 | 7 | 658.91±642.72 | 7 | 167.21±229.99 | 7 | 139.83±141.16 | 7 | | |
| GMCSF | Low | NaN | 0 | 15.63±19.83 | 2 | 2.59±2.81 | 2 | 4.58 | 1 | 12.97 | 1 | NA | NA |
| | High | NaN | 0 | NaN | 0 | NaN | 0 | NaN | 0 | NaN | 0 | | |
| IFNg | Low | NaN | 0 | 6.91±9.48 | 2 | 0.6 | 1 | 1.23 | 1 | 3.42±0.40 | 3 | NA | NA |
| | High | NaN | 0 | NaN | 0 | NaN | 0 | NaN | 0 | NaN | 0 | | |
| IL-10 | Low | NaN | 0 | 61.57 | 1 | NaN | 0 | 76.6 | 1 | 17.96±3.31 | 2 | 0.092 | 0.046 |
| | High | NaN | 0 | 65.75±48.44 | 2 | 50.1 | 1 | NaN | 0 | 290.1 | 1 | | |
| IL-12 p40 | Low | NaN | 0 | NaN | 0 | NaN | 0 | NaN | 0 | NaN | 0 | NA | NA |
| | High | NaN | 0 | NaN | 0 | NaN | 0 | NaN | 0 | NaN | 0 | | |
| IL-12 p70 | Low | NaN | 0 | 3.45±2.76 | 2 | 1.07±0.55 | 3 | 1.08±0.46 | 4 | 2.71±1.44 | 4 | 0.254 | 0.686 |
| | High | NaN | 0 | 14.87±22.51 | 4 | 3.58±5.28 | 3 | 2.63±3.21 | 3 | 1.88±1.02 | 4 | | |
| IL-13 | Low | NaN | 0 | 168.60±222.70 | 2 | 10.92±4.71 | 3 | 17.02±12.06 | 3 | 68.32±31.87 | 2 | 0.257 | 0.558 |
| | High | NaN | 0 | 28.4 | 1 | 8.24±8.09 | 2 | 18.84±15.44 | 3 | 8.60±8.59 | 2 | | |
| IL-15 | Low | NaN | 0 | 39.08±49.05 | 2 | 3.99±2.29 | 3 | 9.90±12.05 | 5 | 14.01±15.90 | 5 | 0.576 | 0.452 |
| | High | NaN | 0 | 14.92±14.68 | 4 | 12.91±13.49 | 3 | 6.44±5.41 | 3 | 41.35±66.66 | 4 | | |
| IL-17 | Low | NaN | 0 | NaN | 0 | 4.6 | 1 | 3.6 | 1 | 2 | 1 | NA | NA |
| | High | NaN | 0 | NaN | 0 | 0.2 | 1 | NaN | 0 | NaN | 0 | | |
| IL-1a | Low | 138.10±145.98 | 3 | 45.66±45.97 | 4 | 114.63±173.67 | 6 | 189.62±242.31 | 6 | 107.59±178.04 | 6 | 0.321 | 0.679 |
| | High | 66.45±0.64 | 2 | 106.03±104.00 | 6 | 92.42±130.45 | 7 | 54.93±50.20 | 4 | 68.54±44.80 | 6 | | |

| | | | | | | | | | | | | | |
|---|---|---|---|---|---|---|---|---|---|---|---|---|---|
| IL-1b | Low | 5.40±2.83 | 2 | 11.13±16.10 | 5 | 5.64±6.71 | 7 | 30.40±68.17 | 7 | 13.90±14.11 | 5 | 0.367 | 0.683 |
| | High | 4.47±4.46 | 4 | 34.72±54.96 | 6 | 30.20±47.33 | 7 | 5.56±6.27 | 7 | 222.13±576.02 | 7 | | |
| IL-2 | Low | NaN | 0 | 73.76±95.66 | 3 | 27.09±20.95 | 5 | 59.15±62.78 | 4 | 54.50±75.54 | 4 | 0.603 | 0.666 |
| | High | NaN | 0 | 35.86±19.00 | 3 | 58.14±74.86 | 3 | 43.71±38.30 | 3 | 25.35±32.29 | 4 | | |
| IL-3 | Low | NaN | 0 | 4.36±6.80 | 3 | 0.44±0.29 | 3 | 0.47±0.31 | 3 | 1.58±1.45 | 3 | 0.524 | 0.796 |
| | High | NaN | 0 | 1.51 | 1 | 0.73±0.94 | 3 | 1.23±1.27 | 3 | 0.72±0.66 | 3 | | |
| IL-4 | Low | NaN | 0 | 16.32±19.80 | 2 | 2.41±1.26 | 3 | 2.70±1.26 | 3 | 7.25±5.94 | 3 | 0.956 | 0.38 |
| | High | NaN | 0 | 7.22±2.28 | 3 | 8.20±12.20 | 3 | 7.06±7.38 | 3 | 3.60±2.71 | 3 | | |
| IL-5 | Low | 77.45±87.47 | 2 | 41.66±59.77 | 4 | 46.82±41.59 | 7 | 30.60±46.08 | 7 | 26.08±20.48 | 5 | 0.843 | 0.75 |
| | High | 36.20±41.86 | 2 | 39.75±36.86 | 5 | 67.98±70.05 | 7 | 20.01±12.98 | 7 | 16.37±13.49 | 5 | | |
| IL-6 | Low | 81 | 1 | 83.38±81.97 | 4 | 176.26±155.61 | 7 | 38.57±44.00 | 7 | 39.21±34.06 | 6 | 0.794 | 0.833 |
| | High | 29.17±16.00 | 3 | 120.23±56.85 | 7 | 257.82±343.91 | 7 | 18.19±17.04 | 7 | 16.87±13.07 | 7 | | |
| IL-7 | Low | NaN | 0 | 349.91±478.20 | 2 | 10.54 | 1 | 14.85±14.67 | 3 | 76.72±4.06 | 2 | 0.418 | 0.51 |
| | High | NaN | 0 | 14.17±8.30 | 2 | 5.24±2.50 | 2 | NaN | 0 | 44.5 | 1 | | |
| IL-9 | Low | 499.40±452.50 | 3 | 123.85±100.35 | 4 | 226.98±345.96 | 7 | 465.52±749.98 | 6 | 345.69±379.55 | 4 | 0.447 | 0.469 |
| | High | 352.10±231.46 | 5 | 511.36±627.91 | 5 | 644.97±701.69 | 5 | 247.22±284.31 | 6 | 433.01±560.80 | 6 | | |
| IP-10 | Low | 478.13±311.12 | 3 | 202.15±107.49 | 4 | 160.29±146.80 | 7 | 227.82±218.83 | 7 | 202.86±171.44 | 7 | 0.357 | 0.78 |
| | High | 466.25±256.54 | 4 | 375.28±253.91 | 6 | 192.47±151.02 | 7 | 189.38±146.79 | 7 | 252.14±232.75 | 7 | | |
| KC | Low | 101.17±35.33 | 3 | 129.01±93.24 | 5 | 380.74±312.83 | 7 | 217.14±430.82 | 7 | 53.71±38.64 | 7 | 0.871 | 0.676 |
| | High | 106.78±99.39 | 4 | 235.74±251.10 | 7 | 521.08±540.95 | 7 | 65.49±55.51 | 7 | 38.38±20.53 | 7 | | |
| LIF | Low | 1.2 | 1 | 14.27±21.65 | 3 | 2.07±1.38 | 6 | 33.59±45.46 | 5 | 5.65±8.28 | 4 | 0.34 | 0.399 |
| | High | 2.80±0.42 | 2 | 25.37±34.87 | 5 | 22.51±31.10 | 5 | 7.06±5.87 | 5 | 104.00±192.26 | 4 | | |
| LIX | Low | 1077.60±1189.01 | 3 | 762.06±577.61 | 4 | 1152.05±1242.29 | 5 | 2365.73±3160.95 | 7 | 1765.18±2716.32 | 5 | 0.105 | 0.729 |
| | High | 6002.55±4901.29 | 4 | 1542.14±1873.54 | 7 | 5959.16±10137.91 | 7 | 3240.94±5325.48 | 6 | 2794.10±3956.37 | 7 | | |
| MCP-1 | Low | 63.2 | 1 | 105.56±131.88 | 5 | 55.05±44.21 | 7 | 149.70±251.16 | 7 | 97.30±159.52 | 6 | 0.275 | 0.302 |

| | | | | | | | | | | | | | |
|---|---|---|---|---|---|---|---|---|---|---|---|---|---|
| | High | 103.80±107.62 | 2 | 212.13±285.60 | 7 | 136.66±217.59 | 7 | 50.65±49.15 | 6 | 872.83±1667.27 | 4 | | |
| MCSF | Low | 5.4 | 1 | 7.20±9.05 | 2 | 12.38±24.18 | 7 | 2.59±1.98 | 4 | 4.37±4.54 | 4 | 0.518 | 0.898 |
| | High | 6.45±7.57 | 2 | 10.82±17.15 | 3 | 4.27±5.88 | 6 | 3.77±3.81 | 3 | 1.32±1.41 | 5 | | |
| MIG | Low | 226.73±155.87 | 3 | 132.70±51.48 | 2 | 73.83±54.01 | 4 | 184.07±36.52 | 4 | 173.22±190.62 | 4 | 0.271 | 0.762 |
| | High | 296.45±276.56 | 4 | 215.93±60.60 | 4 | 117.00±83.92 | 4 | 124.38±114.44 | 4 | 319.48±242.89 | 4 | | |
| MIP-1a | Low | 59.8 | 1 | 21.77±18.42 | 2 | 283.84±560.35 | 5 | 59.85±72.81 | 6 | 103.42±97.71 | 3 | 0.74 | 0.944 |
| | High | 116.20±76.79 | 2 | 115.12±121.52 | 4 | 169.45±164.28 | 4 | 56.83±55.72 | 4 | 57.75±27.26 | 4 | | |
| MIP-1b | Low | NaN | 0 | 2.31±2.73 | 3 | 3.95±6.13 | 3 | 1.74±1.40 | 3 | 13.45±23.39 | 4 | 0.268 | 0.672 |
| | High | NaN | 0 | 1.70±2.09 | 3 | 0.43±0.21 | 3 | 2.01±2.08 | 3 | 1.17±0.91 | 3 | | |
| MIP-2 | Low | 3.4 | 1 | 23.17±19.04 | 4 | 105.52±246.63 | 7 | 27.88±45.29 | 6 | 61.69±122.20 | 6 | 0.453 | 0.845 |
| | High | 35.30±11.96 | 3 | 52.70±96.33 | 6 | 27.43±23.61 | 5 | 13.66±12.67 | 3 | 15.31±11.47 | 5 | | |
| RANTES | Low | NaN | 0 | 36.58±23.02 | 2 | 9.20±7.62 | 2 | 10.40±4.67 | 3 | 20.26±14.53 | 3 | 0.633 | 0.909 |
| | High | NaN | 0 | 45.8 | 1 | 3.36 | 1 | 18.62±16.80 | 3 | 26.93 | 1 | | |
| TNFa | Low | 3.7 | 1 | 22.65±24.52 | 2 | 3.79±2.79 | 4 | 21.07±34.10 | 4 | 15.38±11.48 | 4 | 0.277 | 0.706 |
| | High | NaN | 0 | 49.58±56.18 | 2 | 27.57±37.66 | 2 | 5.48±7.97 | 3 | 130.58±255.49 | 4 | | |
| VEGF | Low | 1 | 1 | 21.25±20.31 | 3 | 58.83±122.66 | 7 | 7.95±8.24 | 7 | 11.50±12.78 | 6 | 0.825 | 0.998 |
| | High | 1.25±0.21 | 2 | 35.87±54.31 | 6 | 53.16±130.83 | 7 | 17.09±15.45 | 4 | 11.57±10.26 | 5 | | |

**Table S4. Flow Cytometry Antibody Information.**

List of surface markers, fluorophore conjugates, clones, suppliers, and working dilutions used for staining murine immune cell populations. All antibodies were used at the specified dilutions, and Fc receptors were blocked with anti-CD16/32 (clone 93) at 4 µg per 2 million cells to prevent nonspecific binding.

| Target | Conjugate | Clone | Supplier | Antibody Dilution |
|---|---|---|---|---|
| **CD45** | BV 605 | 30-F11 | BioLegend | 1:800 |
| **CD11b** | APC-Cy7 | M1/70 | BD Biosciences | 1:200 |
| **CD11c** | BUV 496 | HL3 | BD Biosciences | 1:200 |
| **Ly-6G** | PE | 1A8 | BioLegend | 1:200 |
| **Ly-6C** | PE-Cy7 | HK1.4 | ThermoFisher | 1:200 |
| **F4/80** | BV 421 | BM8 | BioLegend | 1:200 |
| **CD206** | AF 647 | MR5D3 | Bio-Rad | 1:200 |
| **CD3** | AF 700 | 17A2 | ThermoFisher | 1:200 |
| **CD8** | BV 711 | 53-6.7 | BD Biosciences | 1:200 |
| **CD4** | BV 650 | RM4-5 | BioLegend | 1:200 |
| **CD25** | FITC | 7D4 | BD Biosciences | 1:100 |
| **CD19** | BV480 | 1D3 | BD Biosciences | 1:200 |
| **CD16/32** | Unconjugated | 93 | ThermoFisher | 4 µg / 2 million cells |

**Table S5. Flow Cytometry Cell Type Surface Markers.**

Immune cell populations were identified from $CD45^{+}$ cells and subdivided into myeloid ($CD11b^{+}$) and lymphoid ($CD11b^{-}$) lineages. Subsets were classified using the cell surface markers below.

| | Cell Types | Surface Markers |
|---|---|---|
| **Myeloid** (CD45+ CD11b+) | Monoctyes | $Ly6G^{-}$ $Ly6C^{+}$ |
| | Monocytes (M1) | $Ly6G^{-}$ $Ly6C^{Hi}$ |
| | Monoctyes (M2) | $Ly6G^{-}$ $Ly6C^{Lo}$ |
| | Macrophage (M1) | $Ly6G^{-}$ $Ly6C^{+}$ $F4/80^{+}$ |
| | Macrophage (M2) | $Ly6G^{-}$ $F4/80^{+}$ $CD206^{+}$ |
| | Dendritic cells | $Ly6G^{-}$ $CD11c^{+}$ |
| | Neutrophils | $Ly6G^{+}$ |
| **Lymphoid** (CD45+ CD11b-) | T cells | CD3+ |
| | T cells (Cytotoxic) | $CD3^{+}$ $CD8^{+}$ |
| | T cells (Helper) | $CD3^{+}$ $CD4^{+}$ |
| | T cells (Treg) | $CD3^{+}$ $CD4^{+}$ $CD25^{+}$ |
| | B cells | $CD19^{+}$ |
